# Most azole antifungal resistance mutations in the drug target provide cross-resistance and carry no intrinsic fitness cost

**DOI:** 10.1101/2023.12.13.571438

**Authors:** Camille Bédard, Isabelle Gagnon-Arsenault, Jonathan Boisvert, Samuel Plante, Alexandre K. Dubé, Alicia Pageau, Anna Fijarczyk, Jehoshua Sharma, Laetitia Maroc, Rebecca S. Shapiro, Christian R. Landry

**Affiliations:** Institut de Biologie Intégrative et des Systèmes (IBIS), Université Laval, G1V 0A6, Québec, Canada; Département de Biochimie, de Microbiologie et de Bio-informatique, Faculté des Sciences et de Génie, Université Laval, G1V 0A6, Québec, Canada; Département de Biologie, Faculté des Sciences et de Génie, Université Laval, G1V 0A6, Québec, Canada; PROTEO, Le regroupement québécois de recherche sur la fonction, l’ingénierie et les applications des protéines, Université Laval, G1V 0A6, Québec, Canada; Centre de Recherche sur les Données Massives (CRDM), Université Laval, G1V 0A6, Québec, Canada; Department of Molecular and Cellular Biology, University of Guelph, N1G 2W1, Guelph, Canada

## Abstract

Azole antifungals are among the most frequently used drugs to treat fungal infections. Amino acid substitutions in and around the binding site of the azole target Erg11 (Cyp51) are a common resistance mechanism in pathogenic yeasts such as *Candida albicans*. How many and which mutations confer resistance, and at what cost, is however largely unknown. Here, we measure the impact of nearly 4,000 amino acid variants of the Erg11 ligand binding pocket on the susceptibility to six medical azoles. We find that a large fraction of amino acid substitutions lead to resistance (33%), most resistance mutations confer cross-resistance to two or more azoles (88%) and most importantly, only a handful of resistance mutations show a significant fitness cost in the absence of drug (9%). Our results reveal that resistance to azoles can arise through a large set of mutations and this will likely lead to azole pan-resistance, with very little evolutionary compromise. Such a resource will help inform treatment choices in clinical settings and guide the development of new drugs.

## Introduction

Antimicrobial resistance is a global development and health threat^1^. Drug resistance in fungi is particularly challenging because there is a very limited set of treatments to choose from, which means that treatment options are quickly exhausted when resistance mutations occur in patients^2^. Contrary to antibiotic resistance, antifungal resistance is not driven by the acquisition of genes but rather through mutations that occur *de novo* in the genome^3^. Accordingly, the mechanisms of resistance can be very diverse and difficult to detect and predict.

*Candida* spp. are the main culprits of human fungal diseases, causing infections ranging from superficial to invasive. Opportunistic pathogenic yeasts are part of the microbiota of healthy people and cause invasive diseases when the host immune system is compromised^4^. Among the species involved, *Candida albicans* is the most common cause of candidiasis^5^ and it can lead up to 70% mortality when infections become invasive^4^. For these reasons and others, *C. albicans* has been classified as critical in the World Health Organization fungal pathogens priority list^6^.

*Candida* infections are often treated with azoles^4,7,8^. Azoles target the cytochrome P450 lanosterol 14-alpha-demethylase (Erg11 in yeast and Cyp51 in molds), an important enzyme in the ergosterol biosynthesis pathway. When Erg11 is inhibited, ergosterol biosynthesis is blocked, which leads to the accumulation of a toxic sterol intermediate^9^. The evolution of resistance to azoles jeopardizes our ability to treat infected patients^10^. The widespread use of these molecules for treatment or as prophylaxis has led to the emergence of less susceptible isolates^11^. Amino acid substitutions in Erg11 are a common resistance mechanism in most *Candida* spp.^12,13^ Resistance mutations can occur during treatment^2^ and can lead to resistance against multiple azoles simultaneously, a phenomenon known as cross-resistance^3^. Resistance to azoles is not restricted to yeast infections as these drugs are also used to treat other fungal infections such as the ones caused by *Aspergillus spp*. The fact that azoles are also used in agriculture may further exacerbate the resistance problem as resistance mutations are selected in nature, even before fungi infect humans^14–16^.

The evolution of antimicrobial resistance results from mutations that confer an advantage during drug treatment. An important factor that dictates the fate of resistance mutations is whether they reduce fitness in the absence of drugs, for instance following treatment interruption or due to spatial and temporal concentration heterogeneities^17,18^. Mutations with no or low costs are more likely to persist in populations in the absence of drugs, which can influence resistance management strategies^19^. Because antifungal drugs target essential genes or cellular processes, one could expect that resistance mutations would typically alter important protein functions and thus lead to significant fitness costs. Indeed, fitness tradeoffs linked to different resistance mechanisms have been characterized for all clinical antifungal classes^20–23^. Since lanosterol enters the catalytic pocket in Erg11 where azoles also do, one could expect a strong tradeoff for azole resistance mutations. Amino acid substitutions altering azole binding would also impair lanosterol binding and demethylation. However, studies have shown that certain *ERG11* resistance mutations are not costly. For example, the amino acid substitution G464S, which is often observed in clinical resistant isolates of *C. albicans,* does not lead to any growth defect in the absence of drugs, at least in the laboratory^24^. The same has been observed for substitutions G54W and M220K in Cyp51A of *Aspergillus fumigatus*^25^. However, these observations could be biased since resistance mutations that are not costly are more likely to be sampled from patients and isolated in the laboratory. Accordingly, we do not have a clear picture of the extent of fitness cost to azole resistance due to mutations in *ERG11*.

The number of antifungal resistance mutations inventoried in the literature is relatively small compared to all possible antimicrobial resistance mutations and, most importantly, only rarely has the causal link between the mutation and its resistance profile been experimentally validated^12^. Therefore, our knowledge about resistance mutations is insufficient to confidently predict if an amino acid change in Erg11 is going to lead to resistance or not, and what would be the cost associated with these mutations. There is a need for a comprehensive assessment of resistance mutations. Specifically, a better and more comprehensive understanding of the molecular mechanisms underlying resistance, cross-resistance and the associated tradeoffs would enable a better interpretation of the functional significance of *ERG11* genetic variants in *Candida* spp. genomes. Genomic tools could be useful to detect and monitor resistance mutations in a pathogen infecting a patient in real time throughout treatment, allowing for the adjustment of therapeutic use prior to treatment failure^26–28^. Another benefit from such knowledge would be to potentially modify current drugs to overcome or prevent resistance. For instance, a complete map of the resistance mutations in the binding pocket of a target would enable the rational design of drugs that specifically bind the resistant variants or that enter the pocket independently of the mutated residues^29^. Knowledge of resistance mutations would enable the development of effective drugs against resistant isolates without the need to discover completely new molecules or new target genes, which is a limiting factor for fungi because of their shared genes with humans. Finally, there is a large body of research currently attempting to find molecules that would potentiate existing antifungals or re-sensitize fungi to the drugs^30^. A better understanding of resistance mutations and their associated costs would allow us to strategically and rationally identify potentiators. The development of potentiators involves directing efforts towards pathways indirectly implicated in resistance, which are rendered vulnerable by the tradeoffs coming with resistance mutations. For all of these reasons, we aimed to comprehensively identify mutations imparting resistance to azoles, in and around their binding pocket in the enzyme Erg11 from *C. albicans*.

Pathogenic fungi are not easy to manipulate genetically. Methods have been developed and improved^31^ but it is still challenging to engineer a large number of trackable mutants. However, studies have shown that *ERG11* from pathogens can be expressed in the model yeast *Saccharomyces cerevisiae* and complement *ScERG11*’s loss of function, enabling the assessment of the function and resistance of mutations in this model^32–38^. We thus constructed the largest set of Erg11 mutations to date, with nearly 4,000 mutants of *C. albicans* expressed in *S. cerevisiae* engineered to act as a reporter for Erg11 activity. We competed all variants in parallel in the presence of six different azoles to measure resistance to individual drugs and cross-resistance, and in the absence of azole to assess the cost of mutations on the protein function. Our experiments allowed a thorough characterization of the protein variants leading to an extensive atlas of experimentally validated resistance mutations. Further analysis revealed that most azole resistance mutations in *ERG11* provide cross-resistance to azoles and carry no fitness cost. In addition to recovering previously reported high-confidence *CaERG11* resistance mutations, we validated one novel resistance mutation in *C. albicans* by editing *ERG11* directly in its genome and confirmed that it did not confer any fitness cost.

## Results

### Heterologous expression system

As *ERG11* is an essential gene in *S. cerevisiae,* we engineered a yeast strain to make *ScERG11’s* expression repressible so it can be switched off on demand. We constructed a yeast strain with the *ERG11* promoter replaced with a doxycycline repressible one (*ScERG11*-DOX)^39^. This promoter is fully activated in the absence of doxycycline. In the presence of doxycycline, *ScERG11* is repressed enough to completely prevent growth. The expression of *CaERG11* from a plasmid rescues *ScERG11*-DOX growth in doxycycline medium, validating that *CaERG11* complements the *ScERG11* function in this context (Fig. 1a, Extended Data Fig. 1).

**Fig. 1.**
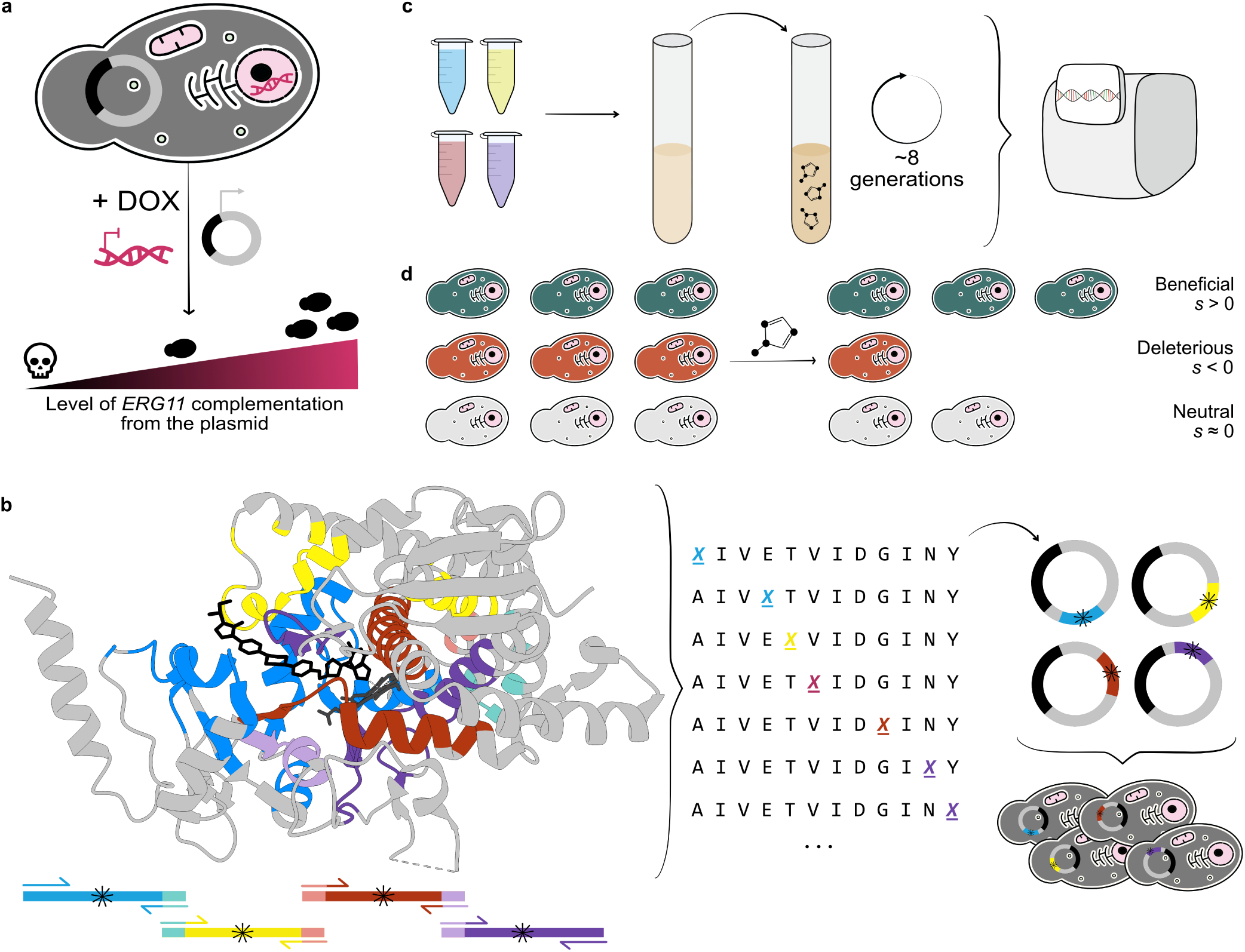
Experimental system for the systematic identification of azole resistance mutations in their drug target Erg11. **a,** *CaERG11* heterologous expression system in *S. cerevisiae* (*ScERG11*-DOX). We replaced the endogenous promoter of *ScERG11* with a doxycycline (DOX on the figure) repressible promoter that is fully active in the absence of doxycycline. When doxycycline is added to the culture medium, *ScERG11* is repressed, which prevents growth. A plasmid expressing *CaERG11* rescues *ScERG11*-DOX growth. **b,** Tertiary structure of CaErg11 bound to itraconazole (Protein Data Bank (PDB): 5V5Z) and mutagenesis. Colored residues were systematically mutated to all alternative amino acids. The library was split into four overlapping fragments to allow sequencing with short reads. The variant library was cloned into plasmids containing the native *ERG11* promoter and terminator of *S. cerevisiae* and transformed into the strain *ScERG11*-DOX. **c,** Pooled competition assays. The yeast library was pooled by fragments and grown without antifungal selection pressure. Then, it was transferred into a medium containing doxycycline and an azole, and cells were left to compete for eight generations. Variant abundance at the beginning and at the end of the competition was estimated by high-throughput sequencing. **d,** Selection coefficient (*s*) to phenotype. Variant abundance before and after the competition was used to calculate selection coefficients. Variants were classified into three categories: beneficial, deleterious and neutral.

### *CaERG11* deep mutational scan

Since most amino acid substitutions shown to confer resistance are found in regions adjacent to the enzyme’s active site^7,40^, we concentrated our efforts on residues most likely to perturb interactions with azoles. We examined the protein structures to find residues close to the ligand binding pocket. All the residues selected are located at less than 12 Å either from the lanosterol, fluconazole, itraconazole and heme molecules in the protein (Extended Data Fig. 2a). This range was chosen to include all well-characterized resistance mutations, including those further away from the ligand binding site such as Y257H (11.80 Å)^37,38,41–44^ (Extended Data Fig. 2b).

The structural analysis led to a selection of 206 positions for the variant library (Fig. 1b, Extended Data Fig. 2c,d, Supplementary data 1). The library was designed to include the wild-type sequence and all 206 residues individually replaced by the 19 other amino acids. We used two codons per amino acid to create two biological replicates, except for methionine and tryptophan that are encoded by a single one (Supplementary data 2). *CaERG11* variants were synthesized and cloned into a plasmid containing the native promoter and terminator of *ScERG11* to ensure that expression regulation is similar to that of the native gene. We divided *CaERG11* into four overlapping sections to have DNA fragments of suitable size for high-throughput sequencing and we pooled the variants according to the fragment where their mutation is located (Fig. 1b). We transformed all plasmids into the *ScERG11-*DOX strain. The final library contains 3,830 amino acid variants (Extended Data Fig. 3).

We competed the variants with and without an azole to assess resistance and the fitness cost of mutations (Fig. 1c, Extended Data Fig. 4). We selected six azoles that are used in clinics. These molecules have more or less different structures. Fluconazole and itraconazole are first-generation triazoles, and their second-generation counterparts, voriconazole and posaconazole have similar chemical structures. Isavuconazole is one of the newest triazole. Clotrimazole is an imidazole widely used to treat superficial infections^45^ (Fig. 2a). The concentration inhibiting around 50% of wild-type growth (IC_50_, Extended Data Fig. 5) was used for the pooled competition assays in each case.

**Fig. 2.**
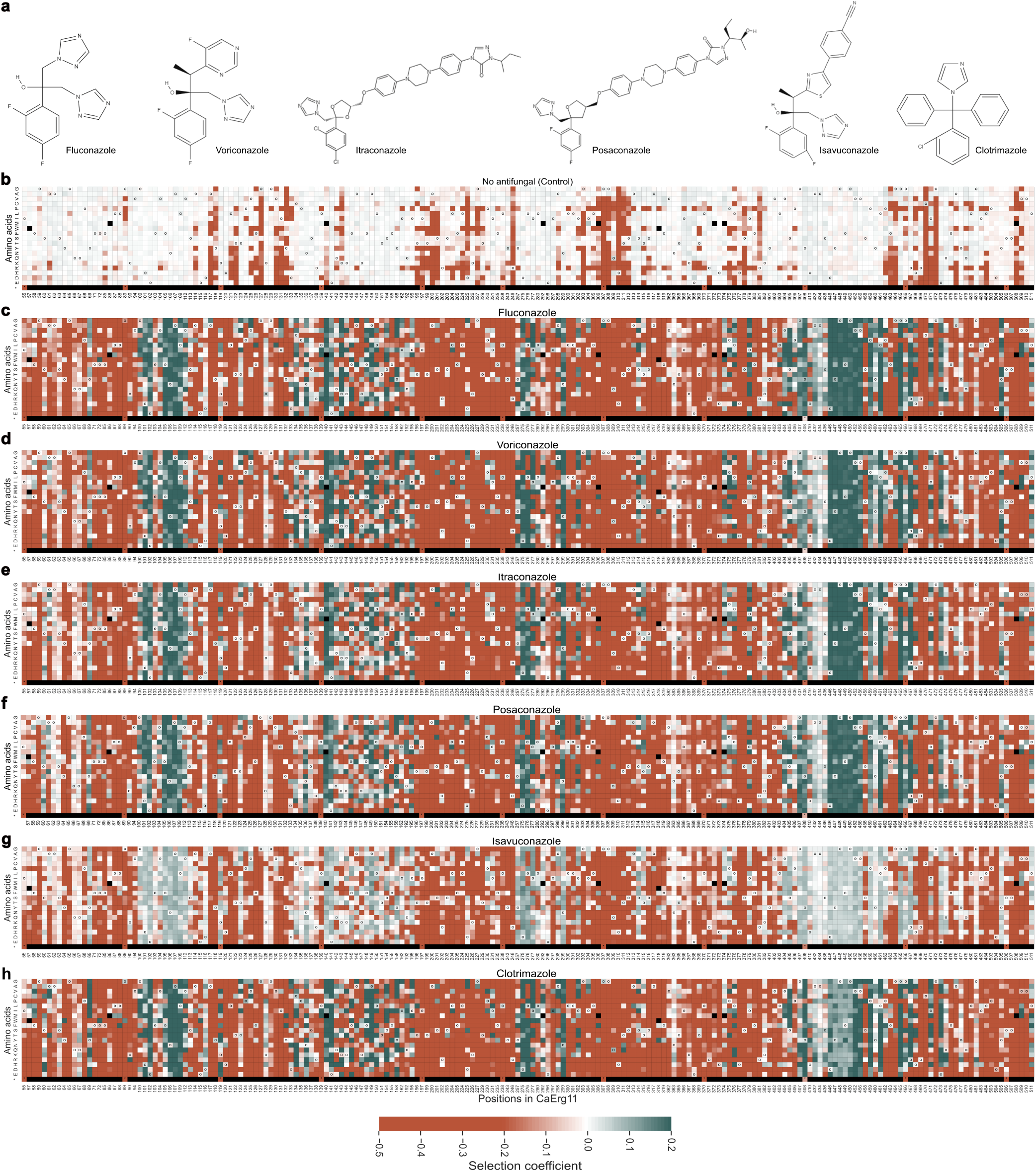
The resistance landscape of *C. albicans* Erg11 to six clinical azoles. **a,** Chemical structures of the azoles used. Structures were retrieved from PubChem^46^. **b,c,d,e,f,g,h,** Selection coefficients (*s*) of CaErg11 variants in control condition (**b**) and in fluconazole (**c**), voriconazole (**d**), itraconazole (**e**), posaconazole (**f**), isavuconazole (**g**) and clotrimazole (**h**). *s* > 0 (green) indicates higher fitness than the wild-type and *s* < 0 (orange) indicates lower fitness than the wild-type. “o” represents wild-type amino acids and * represents stop codons (stop codons were introduced as controls at every 20 residues). n = 3,830 amino acid variants. Each value is the median of two synonymous codons per amino acid (in most cases), and triplicate experimental measurements. Black cells represent missing values (wild-type amino acids tryptophan and methionine do not have synonymous codons).

We measured variant frequencies by high-throughput sequencing at the start and the end of the eight generations of cell competition. Variant sequence coverage was higher than 700x on average per sample at the initial and final time points, providing high-quality data. The difference between the initial and final frequencies was used to calculate a selection coefficient (*s*) of all amino acid variants (Fig. 2b-h). To control for the possibility of mutations occurring in other genes of the *S. cerevisiae* genome and being selected for during the competitions, we included several replicates at the codon and gene fragment levels, and we replicated the cultures. A spontaneous genomic resistance mutation that would occur in one initial cell in the culture would therefore be associated with one codon for a given amino acid and one specific replicate for a given codon. Consequently, the impact of such mutations would be to weaken the correlations among replicates. We conclude that spontaneous mutations in the cultures, if any, are not significantly influencing growth of variants in the pooled assays since the measurements were highly reproducible, with Spearman’s rank correlation coefficient (ρ) for the same codons among replicates ranging from 0.95 to 0.99 and for the same variants in overlapping fragment positions ranging from 0.98 to 1.0 (*P* < 0.0001, Extended Data Fig. 6 and 7). The selection coefficients of synonymous codons for each given amino acid were also strongly correlated (*ρ* between 0.92 and 0.98, *P* < 0.0001, Extended Data Fig. 8). The range of selection coefficients varies among azoles, but they all have similar distribution shapes (Extended Data Fig. 9).

We used the selection coefficient estimates (*s*) of individual amino acid variants to infer their resistance to azoles. We classified the variants into three categories: beneficial, deleterious and neutral (Fig. 1d). We performed a *t-*test to assess whether *s* estimates of missense variants were significantly different from that of the wild-type sequence. We defined variants as beneficial or deleterious if the *p-*value adjusted for false discovery rate (FDR) was lower than 0.01.

### Impact of mutations on the function of Erg11

We first focused on the impact of mutations on growth in the control condition (without azole). Not surprisingly for an essential enzyme, the stop codons included as controls are highly deleterious (bottom row of heatmaps in Fig. 2b-h) and some amino acid substitutions are as deleterious as stop codons. Mutations that completely abolish the enzyme function form clusters along the sequence (Fig. 3a). Positions near residues forming a hydrogen bond with the heme (Y118, Y132, K143, R381, H468 and C470) and near residues participating in proton transport to the heme (D225, H310 and T311) are the most sensitive to amino acid changes^47^. These positions are usually highly conserved among fungal orthologs, which validates their important role in Erg11 canonical enzymatic reaction, more specifically in lanosterol demethylation (Fig. 3b, Extended Data Fig. 10).

**Fig. 3.**
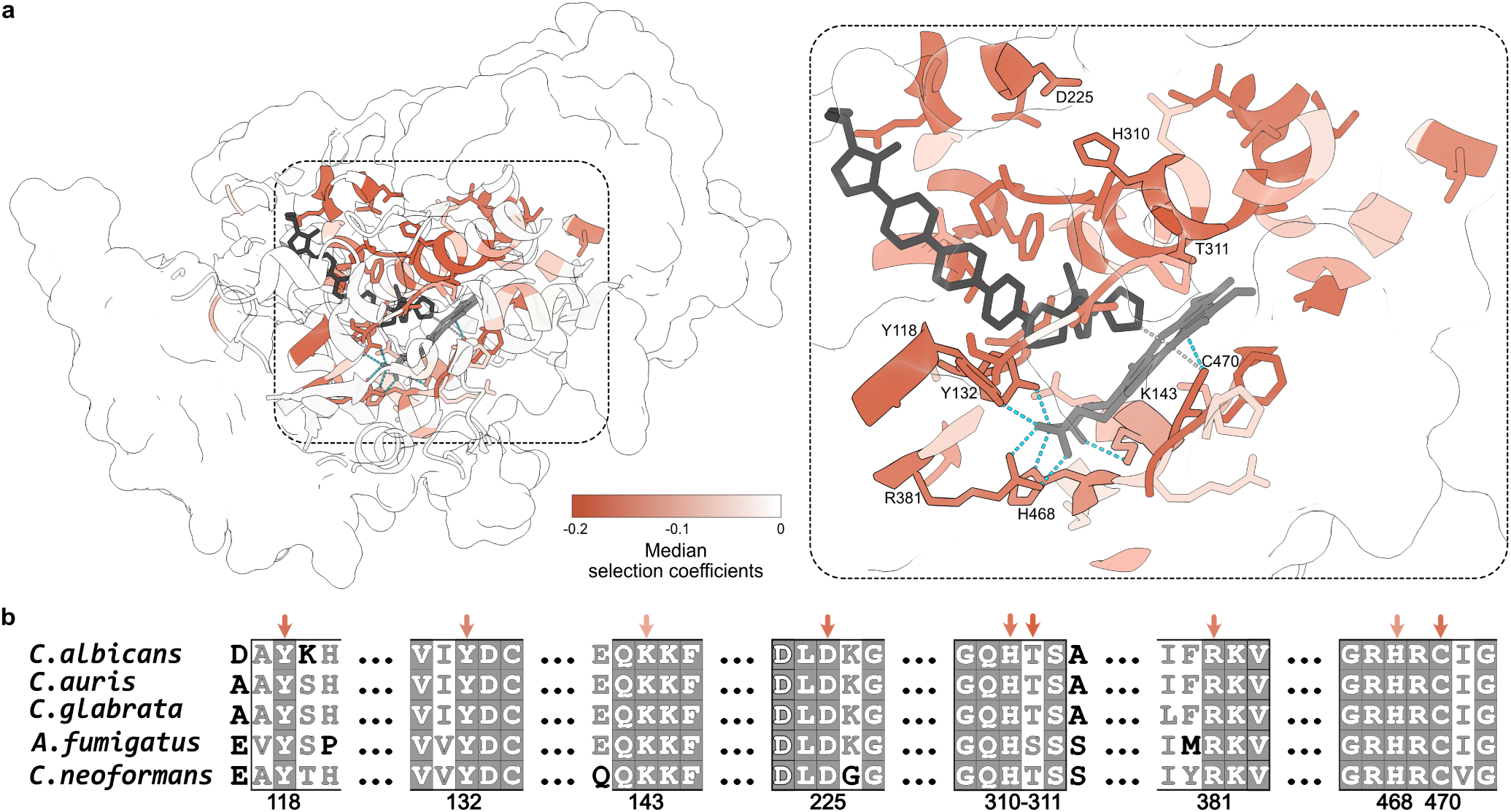
Mutations impairing CaErg11 canonical function are clustered and occur at conserved sites. **a,** Median selection coefficients in the control condition (without azole) mapped on CaErg11 bound to itraconazole (PDB ID : 5V5Z). On the right, only positions with deleterious variants are shown for clarity. Hydrogen bonds are represented as cyan dotted lines, itraconazole is in dark grey and the heme molecule is in light grey. **b,** Multiple sequence alignment of Erg11 for some fungal pathogens listed on the WHO priority list (*Candida albicans, Candida auris, Candida glabrata, Aspergillus fumigatus* and *Cryptococcus neoformans*)^6^. A partial sequence is shown to highlight conserved amino acids interacting with the heme (Y118, Y132, K143, R381, H468 and C470) and participating in proton transport to the heme (D225, H310 and T311). The complete alignment is available in Extended Data Fig. 10 and was produced with ENDscript 2^54^.

To further validate our measurements, we examined the relationship between long-term evolutionary constraints on the enzyme and the impact of mutations on fitness measured here. We estimated the potential impact of mutations using patterns of evolution of individual positions among orthologs as implemented in GEMME^48^. The selection coefficients measured in the control condition show a highly significant positive correlation with the evolution-based predictions (*ρ* = 0.40, *P* < 0.0001). Interestingly, we observed fewer deleterious mutations than what GEMME predicts (Extended Data Fig. 11a). One reason for the discrepancy between the predictions and the measurements could be that there were recent changes in structure or function of Erg11 that failed to be captured by comparing very distant orthologs. The experimental conditions used here also do not reflect all the conditions this enzyme could have been selected to perform in. In addition to altering the enzymatic activity, one other reason why mutations could be deleterious for the function of Erg11 is protein destabilization. We therefore used FoldX^49^ to predict the effects of all amino acid substitutions on CaErg11 stability. We observed a negative correlation (*ρ* = −0.27, *P* < 0.0001) between the change in free energy (ΔΔG) predicted with FoldX and *s*. This correlation suggests that some amino acid substitutions are deleterious because they destabilize the enzyme, but it is likely not the only factor (Extended Data Fig. 11b).

To evaluate if the deleterious effects we measured in *S. cerevisiae* can be generalized to natural *C. albicans* populations, we analyzed 435 genomes from four studies that were publicly available^50–53^. Although polymorphisms are limited because we study a single gene, we identified variable amino acid substitutions overlapping the genomic dataset and our DMS library. We found that the mutations we classified as deleterious in the control condition were much less frequent than the ones we classified as neutral (Extended Data Fig. 12). In addition, the only two mutations we identified as deleterious in the control conditions that are present in the 435 genomes are found in only two isolates and are heterozygous (one isolate with G472R and another with H373Y). The higher frequency of mutations in *C. albicans* that we identified in the *S. cerevisiae* expression system as neutral suggests that they are indeed neutral and it shows that the effects we are measuring can be transposed on natural *Candida albicans* populations.

### A large fraction of mutations lead to resistance, including outside of known hotspots

Through the analysis of selection coefficients, we identified a large number of mutations that significantly increase growth rate compared to the wild-type when cells are exposed to antifungals (*s* > 0, *P* < 0.01). We defined all variants with a *s* > 0 as beneficial when exposed to azoles, and thus resistant, even if resistant variants present a wide range of fitness. We identified 946 resistance amino acid substitutions for fluconazole, 1,044 for voriconazole, 999 for itraconazole, 1,011 for posaconazole, 1,009 for isavuconazole and 882 for clotrimazole, for a total of 5,891 mutation-resistance associations.

It has been proposed that resistance mutations cluster into three hotspots^55^ based on sequences from fluconazole and itraconazole resistant clinical isolates. The first hotspot extends from residue 105 to 165, the second from 266 to 287 and the third from 405 to 488. Since then, these hotspots have been used as a reference in numerous papers to help interpret the impact of mutations. In our experiments, we identified resistant mutants in the three clusters, as well as at positions outside. More precisely, residues with more than 10 variants conferring fluconazole resistance mostly fall into the hotspots (Extended data Fig.10). We determined that 40% of the variants included in the three hotspots (709/1752) conferred fluconazole resistance. To visualize on the 3D protein structure where the residues that most often confer resistance when mutated are located, the medians of the selection coefficients were mapped on CaErg11 structure (Fig. 4a). We found that the variants included in the hotspots form altogether a large cluster. However, if we instead mapped on the structure all positions with at least one resistant variant (maximal values of *s* per residue), we discovered that there are many resistant variants occurring outside the initial clusters (Fig. 4b). We estimated that 11% of the variants located outside the hotspots (237/2078) confer fluconazole resistance, while 60% of those inside the hotspots remain sensitive to azoles (1043/1752). In the light of these results, we believe that resistance hotspots should be used with caution as a guide to interpret the impact of mutations in Erg11 since they could potentially lead to inaccurate predictions of resistance.

**Fig. 4.**
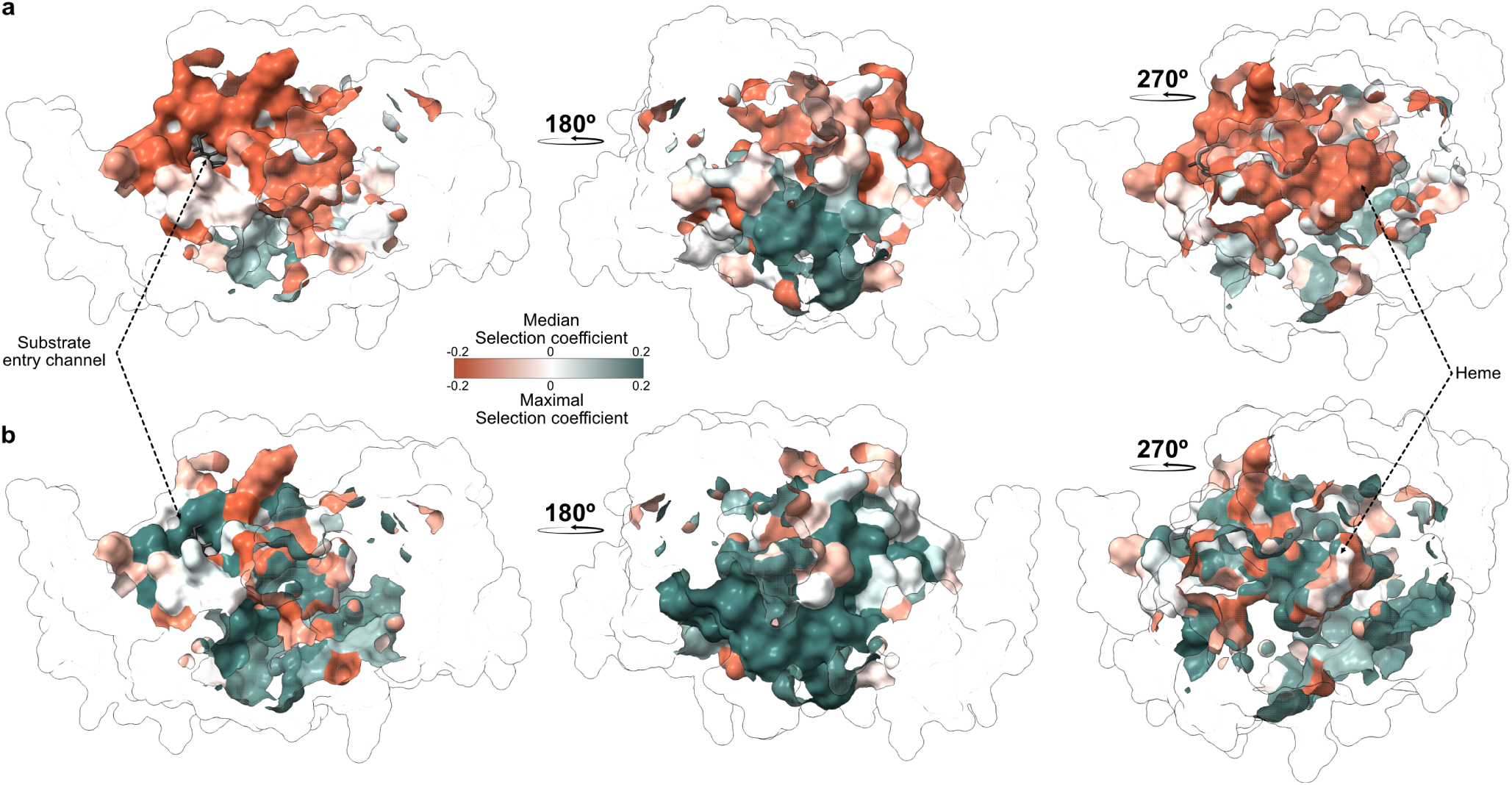
Azole resistance mutations in CaErg11 are scattered across the protein tertiary structure. **a,** CaErg11 (PDB ID 5V5Z) with the median selection coefficients per residue mapped on the structure. On the 180° rotation, a distinct cluster of residues showing strongly positive selection coefficients includes residues located in the three hotspots proposed by Marichal *et al*.^55^ **b,** CaErg11 (PDB ID 5V5Z) with the maximal selection coefficients per residue mapped. We can see that resistance mutations are scattered across the structure and not limited to a single cluster.

We validated the impact of several mutations by reconstructing individual mutants by site-directed mutagenesis of the wild-type *CaERG11* plasmid and by re-assessing the growth of the variants individually. To do so, we constructed 35 variants covering a wide range of *s* in all conditions. The variants were grown for 24h in media containing the azoles tested in the pooled competition assays and in the control condition. The correlation between the growth of the individual mutants and *s* estimated from bulk competition in azoles is strong and highly significant (*ρ* ranging from 0.83 to 0.89, *P* < 0.0001), showing that the high-throughput pooled assays are highly reproducible with individual mutants on smaller scales (Fig. 5a). Most variants in the control condition grew similarly, as anticipated.

**Fig. 5.**
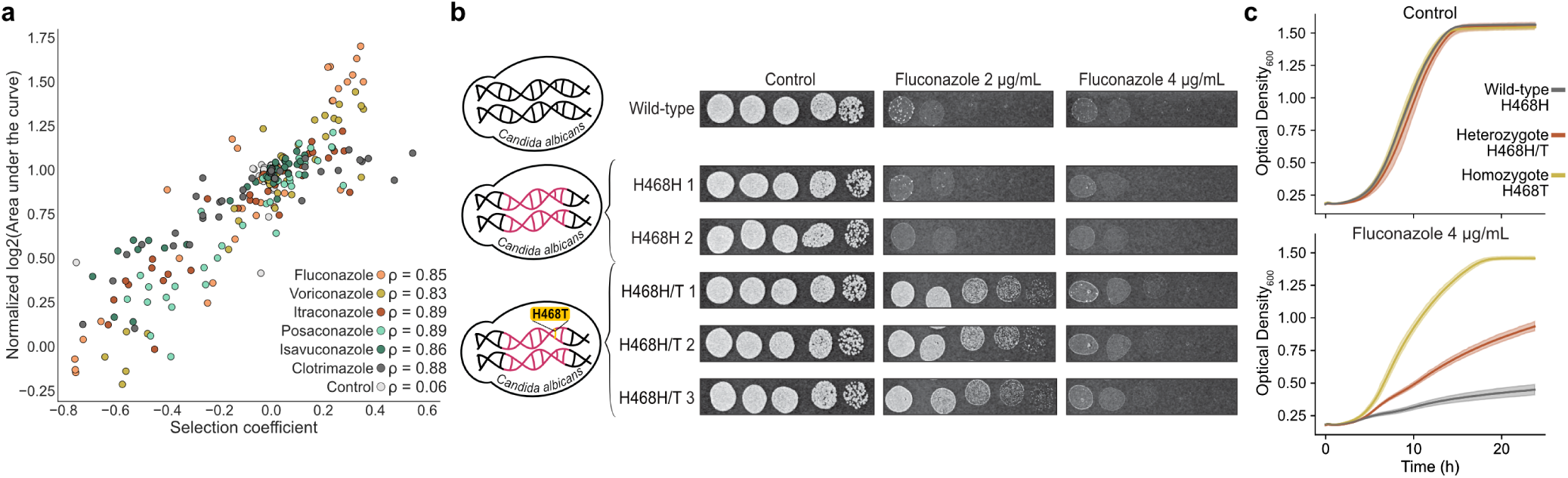
High throughput assays are reproducible at smaller scale in *S. cerevisiae*, and a novel resistance mutation is validated in *C. albicans*. **a,** Validation of 35 variants individually reconstructed. Growth of mutants measured as Log2 of the area under the curve compared with the selection coefficient of the DMS competitions with and without azoles. The values were normalized relative to the wild-type. Colors represent the different conditions. Each point is the average of two replicates. Spearman’s rank correlations are shown, *P* < 0.0001 for all. **b,** Spot assays of Erg11 H468H/T CRISPR-Cas9 genome edited *Candida albicans* compared to Erg11 H468H wild-type CRISPR-Cas9 genome edited controls and SC5314 wild-type strain. Serial 1/5 dilutions of cells starting at an OD_600_ of 1 on YPD medium supplemented or not with fluconazole. **c,** Growth in the control condition and in fluconazole (4 μg/mL) was measured for *Candida albicans ERG11* H468H wild-types (n=2), H468H/T heterozygotes (n=6) and H468T homozygotes obtained through loss of heterozygosity (n=5). All strains presented similar growth in control media and the H468H/T heterozygotes presented an intermediate resistance phenotype in fluconazole in comparison to the wild-types and H468T homozygotes.

### Resistance mutations identified by heterologous expression in *S. cerevisiae* have the same effect in *C. albicans* background

The impact of a mutation may be affected by the genetic background in which it occurs due to genetic interactions^56,57^. Thus, while a mutation in CaErg11 may confer resistance to azoles in our system based on heterologous expression of the gene in *S. cerevisiae*, this effect may not be reproduced if the mutation occurs directly in the *C. albicans* genome background. Therefore, to assess how the results we obtained by heterologous expression of CaErg11 in *S. cerevisiae* background correlate with *C. albicans* clinical isolates, we compared our results with known, experimentally validated resistance mutations in CaErg11. Since *C. albicans* is often subjected to treatments with fluconazole, mutations have been reported and validated experimentally for this particular drug. Amino acid substitutions A114S, T123H, Y132F, Y132H, K143E, K143Q, K143R, F145L, Y257H, S405F, D446E, G448E, F449V, G450E, Y447H, G464S and V456I have all been validated to confer at least a 2-fold decrease in fluconazole susceptibility^37,38,41–44^. All these mutants were classified as resistant in our experiments (*s* = 0.07 to 0.46), except for A114S which was classified as neutral, although it has a slightly positive *s* that is non-significant after correction for multiple testing (*s* = 0.04, *P* = 0.13). Voriconazole and itraconazole are also often used to test for azole susceptibility. Substitutions A114S, T123H, Y132F, Y132H, K143Q, K143R, F145L, Y257H, S405F, G448E, G450E and G464S have been shown to confer a minimum 2-fold decrease in voriconazole and itraconazole susceptibility in at least one study^37,38,41,42^. We classified mutants carrying these substitutions as resistant to voriconazole (*s* = 0.08 to 0.47) and itraconazole (*s* = 0.07 to 0.26), validating that the mutations indeed reduce susceptibility, contrary to what has been reported in some other studies for voriconazole^41^. We provide the selection coefficients and the classification for each CaErg11 variant as a resource (Supplementary data 3).

To further assess if the background significantly affects the impact of mutations in CaErg11, we validated the effects of a newly identified resistance mutation directly in *C. albicans* using CRISPR-Cas9 to edit its genome. The strain we engineered is heterozygous for H468T because editing efficiency was not high enough to edit both gene copies. As expected, the H468H/T strain grew better than the wild-type on fluconazole medium (Fig. 5b).

Resistance mutations we identified in *S. cerevisiae* by expressing a single allele could be recessive. However, reports from the literature and our genome editing experiment suggest that they could be at least partially dominant in a diploid genome. This would be an important aspect of resistance to be selected for, as resistance mutation that would be entirely recessive would not be visible to selection in a diploid species. We note however that loss of heterozygosity (LOH) is a frequent mechanism of resistance in *Candida spp*. such that heterozygous resistance mutations could rapidly become homozygous^58^. To test this hypothesis, we experimentally evolved H468H/T heterozygous edited strains for four days in the presence of increasing concentrations of fluconazole. Using this strategy, we indeed rapidly obtained H468T homozygous strains. Then, we assessed the growth of the H468H wild-type, the H468H/T heterozygotes and the H468T homozygotes in the absence and in the presence of fluconazole. All strains presented similar growth in the control medium. As expected, the H468H/T heterozygotes presented an intermediate resistance phenotype in the presence of fluconazole in comparison to the wild-types and H468T homozygotes (Fig. 5c). Therefore, this mutation provides a greater benefit upon growth in azole antifungals when it is homozygous, and does not incur a detectable fitness cost in the absence of drug.

### Most resistance mutations provide cross-resistance

Despite the fact that azoles have different structures, the patterns of resistance along the sequences appear to be strikingly similar (Fig. 2b-h). Indeed, selection coefficients are strongly correlated among azoles, with Spearman’s rank correlation coefficients ranging from 0.93 to 0.99 (*P* < 0.0001, Extended Data Fig. 13). We found the strongest correlation between itraconazole and posaconazole (ρ = 0.99) followed by isavuconazole and itraconazole or posaconazole (ρ = 0.97). Fluconazole and clotrimazole were the least correlated to others (ρ = 0.93 and 0.94), except for the fluconazole-voriconazole pair (ρ = 0.96). These correlations do not reflect the overall similarities among the molecules. The specificity of azoles may therefore derive from subtle differences in interactions between the molecules and CaErg11.

Overall, 33% (1,268) of the amino acid changes tested confer a significant growth advantage to at least one azole. Of those, 1,117 are resistant to multiple azoles, with 64% (712) being resistant to all of the azoles tested (Fig. 6a). Cross-resistance, whereby a mutation is advantageous in more than one azole, is therefore very common for this class of antifungal. Additionally, one third of the cross-resistance mutations (252/712) are attainable through only one nucleotide change from the wild-type *C. albicans* sequence. This implies that *C. albicans* can very quickly become resistant to all azoles.

**Fig. 6.**
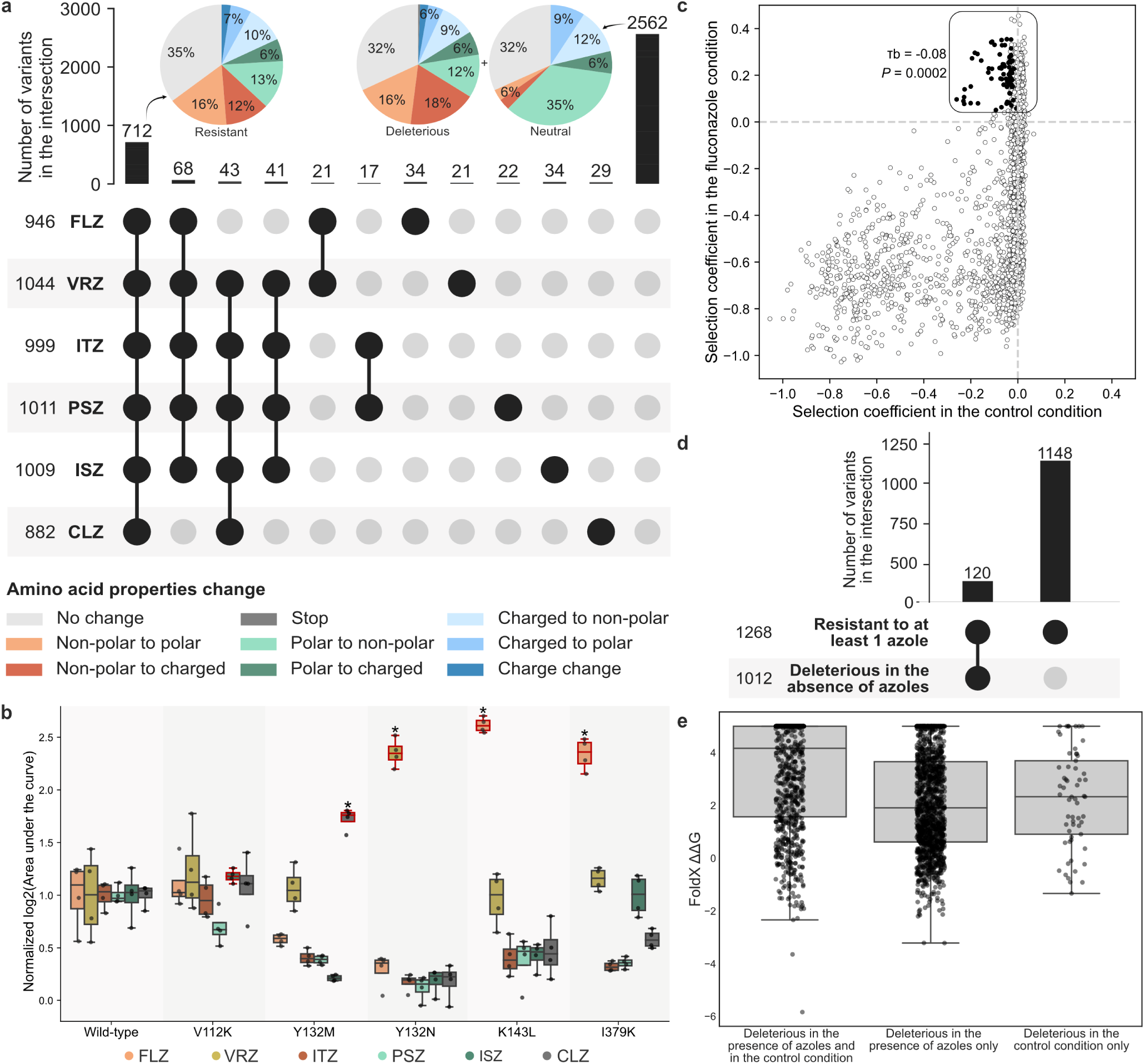
Azole resistance mutations in CaErg11 confer cross-resistance and rarely carry a fitness cost. **a,** Upset plot showing the intersection of azole resistant variants (black circles). The number of resistant variants for each azole is shown as well as the number of variants in the intersections. Only intersections with more than 15 variants are shown for clarity. Pie charts represent the change in amino acid properties for the intersection including variants resistant to all azoles and the intersection including variants neutral or deleterious for all azoles. FLZ: fluconazole, VRZ: voriconazole, ITZ: itraconazole, PSZ: posaconazole, ISZ: isavuconazole and CLZ: clotrimazole. **b,** Log2 of the area under the curve normalized relative to the wild-type for a subset of five variants that showed drug-specific resistance (n=4 replicates)(red overlined boxplots: V112K: isavuconazole, Y132N: voriconazole, Y132M: clotrimazole, K143L: fluconazole and I379K: fluconazole). 4/5 variants grew significantly more than the wild-type for only one azole, showing their specificity (*, Mann–Whitney U test, *P <* 0.05). Colors represent the different conditions and boxes represent the interquartile range. **c,** Selection coefficients of amino acid substitutions in fluconazole pooled competition assays compared with the control condition. Each value is the median of two synonymous codons per amino acid in most cases for the median of triplicate experimental measurements. The dark grey dots represent variants carrying a tradeoff, meaning that the mutation is advantageous in the presence of azole and significantly deleterious in the control condition. Kendall’s tau-b (τb) rank correlation is shown. n = 3,830 amino acid variants. **d,** Upset plot showing the intersection of variants resistant to at least one azole and deleterious in the control condition. The number of variants for every group is shown as well as the number of variants in the intersections. **e,** Change in free energy (ΔΔG) predicted with FoldX for variants categorized as deleterious. ΔΔG values are higher for variants deleterious in both the azole and control conditions (n = 858 variants, median ΔΔG of 4.16) compared to those deleterious only in azoles (n = 1331 variants, median ΔΔG of 1.91) or only in the control condition (n = 61 variants, median ΔΔG of 2.33) (Mann-Whitney U test, *P* < 0.0001). Values higher than five were rounded to five for plotting.

To further rationalize the resistance conferred by certain amino acid substitutions, we examined whether the alteration in amino acid properties was significantly different between the categories of variants. Notably, a dissimilarity in amino acid properties was observed between resistant and neutral variants (chi-squared test, *P* = 0.0003), but not with deleterious variants (chi-squared test, *P* = 0.97) (Fig. 6a). Consequently, the general shift in amino acid properties alone cannot account for the observed resistance.

Even though cross-resistance between azoles seems to be common, we note some interesting exceptions. We categorized 151 variants resistant to only one azole and we could pinpoint specific residues which, when mutated, confer resistance only to a small subset of azoles. For instance, almost all variants at position 505 confer resistance to fluconazole and voriconazole, but none of them confer resistance to clotrimazole. Similarly, more variants at position 464 are resistant to fluconazole and voriconazole compared with the other azoles. Interestingly, G464S is one of the most often reported resistance mutations in the literature^12,42^, and it is the only variant at position 464 resistant to all azoles in our experiment. Among all drug-specific variants, we chose five to ascertain their specificity by reassessing their growth individually. Subsequent validations revealed that 80% (4/5 variants) displayed significantly higher growth than the wild-type in only one condition, confirming drug-specific resistance (Mann–Whitney U test, *P* < 0.05) (Fig. 6b). Therefore, despite the drug-specific variants constituting a small proportion of the resistant variants, their presence is noteworthy and may yield valuable insights for targeted intervention strategies.

### Most resistance mutations carry no significant fitness cost

As Erg11 is a key enzyme in the ergosterol biosynthesis pathway and since its canonical substrate sits in the same binding pocket as azoles, resistance mutations have a strong potential to carry a fitness cost in the absence of drugs. One would therefore expect a tradeoff between resistance and fitness in the absence of azoles. We first tested if there was a negative correlation between the selection coefficient of amino acid substitutions conferring azole resistance and the control condition. A negative correlation would show that in general, resistance mutations negatively alter the enzyme function sufficiently to reduce growth in control conditions. Surprisingly, we only found weak correlations for most azole conditions (Kendall’s tau-b (τb) = −0.08 to 0.09, *P* = 0.24 to 0.00005) (Fig. 6c, Extended Data Fig. 14). This almost absence of correlation shows that many mutations enhance drug resistance without altering protein function. Second, we examined the fraction of resistant variants that are also deleterious in the control condition. Our analysis revealed that only 9% (120) of resistant variants carry a significant fitness cost (Fig. 6d). Positions 140, 143 and 471 are particularly enriched for such tradeoffs, probably because of their interaction or proximity with the heme^47^ (Fig. 3a). Since amino acid substitutions 143E and Q have been reported in clinical isolates^12,38^, our results suggest that tradeoffs between resistance and fitness are relatively rare for Erg11 and that when they occur, they do not prevent mutations from occurring in clinical settings. Either selection is too strong for the tradeoffs to play a significant role or compensatory mutations or physiological mechanisms (e.g. upregulation or downregulation of other genes) reduce the impacts of costly mutations during drug treatment^19,59^.

Increased Erg11 abundance is a known azole resistance mechanism^9^. The larger number of enzyme molecules in the cell compensates for azole inhibition. To examine the possibility that some of the mutations increase CaErg11 abundance, which could lead to resistance without a significant fitness cost, we fused the green fluorescent protein (GFP) to a subset of the CaErg11 variants that we individually reconstructed (n = 17 mutants and n=1 wild-type) and we measured protein levels by flow cytometry (Extended data Fig.15). Among the eight fluconazole-resistant variants assessed, only two (V130L and K143L) showed significantly higher protein levels than the wild-type (Mann-Whitney U test, *P* < 0.05). Increased protein abundance is thus a potential factor but it is not the main factor to explain resistance. Interestingly, the two resistant mutants with increased abundance are carrying a fitness cost, suggesting that higher protein abundance also carries negative consequences. Furthermore, out of the four variants with a higher susceptibility to fluconazole, two (K147C and L480K) presented significantly different protein levels than the wild-type (Mann-Whitney U test, P < 0.05). L480K had significantly lower protein levels while K147C had significantly higher protein levels. For L480K, its lower abundance could be enough to sustain wild type-level growth in control conditions, but when an azole is present, its lower abundance combined with inhibition leads to decreased growth. The higher fluconazole susceptibility of K147C, despite increased protein levels, could potentially be explained by an increased affinity to azoles. These results suggest that protein destabilization is not the only cause of the deleterious effects observed but that it can play a role.

Despite the fact that few amino acid substitutions significantly impaired CaErg11 function in the control condition, a larger number were highly harmful in the presence of azoles (Fig. 2b-h) while having a mild or no effect in control conditions. Specifically, 26% (1,012 variants) were found to be deleterious in the control condition, while this percentage increased to 58% (2,204 variants) in azoles. It is likely that certain mutations slightly diminish the canonical enzymatic reaction, without exerting a substantial impact on fitness under normal conditions. However, their effects would be amplified in the presence of drugs that diminish the enzyme function below a level that is required for normal growth, resulting in an increased susceptibility to azoles. We observed that the changes in free energy (ΔΔG) predicted with FoldX^49^ are larger for variants deleterious in both the azole and control conditions (n = 858 variants, median ΔΔG of 4.16) compared to those deleterious only in azoles (n = 1331 variants, median ΔΔG of 1.91) (Fig. 6e, Mann-Whitney U test, *P* < 0.0001). This is consistent with a slight destabilization by the mutations that reduce enzyme activity or abundance but that is visible only when the drug further inhibits its function. Highly destabilizing mutations on the other hand would already have an effect without azoles. In a similar way, mutations deleterious only in the control and not in azole conditions have significantly lower predicted ΔΔG in comparison with mutations that are deleterious in all conditions, since highly destabilizing mutations would be deleterious both in control and in the presence of azoles (n = 61 variants, median ΔΔG of 2.33) (Fig. 6e, Mann-Whitney U test, *P* < 0.0001). Other drugs that could destabilize Erg11 or reduce its abundance could therefore be interesting leads for sensitizing fungi to azoles.

It is important to note that the experimental conditions used here do not necessarily reflect the selective pressure of the natural or clinical environments. Therefore, the proportion of tradeoff mutations could be more important if one considered other potential environments that could make the function of Erg11 even more critical or that could further enhance the impact of resistance amino acid substitutions, for instance elevated temperatures^17,60^. Also, it is possible that the fitness cost of some resistance mutations that we measured here is larger in *C. albicans* than it is in *S. cerevisiae,* although we consider that unlikely since Erg11 is essential for *S. cerevisiae* but not for *C. albicans* according to some reports^61,62^. We therefore think that our assays are conservative. Furthermore, a study in *Candida parapsilosis* also noted the absence of a cost for a resistance mutation in *ERG11* in a *Galleria mellonella* model of infection^63^.

## Discussion

Infections caused by fungal pathogens are a serious public health problem^64^. The emergence of fungi resistant to antifungals threatens our ability to treat infections^10,11,65^. *C. albicans* is the most frequent cause of fungal infections in humans, which are routinely treated with azole drugs that inhibit the enzyme cytochrome P450 lanosterol 14-alpha-demethylase (Erg11). Despite the key importance of this species and of Erg11 in clinical microbiology, our picture of the drug resistance mechanisms conferred by mutations in *ERG11* is incomplete. Here, using a deep mutational scanning approach, we created a library of nearly 4,000 variants of *C. albicans* Erg11. The library was used to characterize resistance mutations to six medical azoles, leading to the mapping of more than 25,000 phenotypes to protein sequence genotypes. For each drug, an average of 1,000 variants were found to confer resistance, which led to nearly 6,000 experimentally validated drug-resistance associations. More than 50% of the resistant variants are resistant to all of the azoles tested and less than 9% of them lead to a fitness defect in the absence of drugs. The systematic characterization of Ca*ERG11* variants allowed a better understanding of cross-resistance and tradeoff which we hope will impact regulations linked to azole usage and help develop new drugs. Additionally, the creation of an extensive catalog of Ca*ERG11* mutations leading to azole resistance will improve sequence information interpretation and could help treatment choice in clinical settings. In this study, we targeted the main gene involved in azole resistance and tested the most frequently used antifungals. As many other molecular mechanisms lead to antifungal resistance, using a similar approach as this one or as that of Desprès et al.^21^ and Rouleau et al.^66^ on other fungal genes will be necessary to profile drug resistance mutations comprehensively and ultimately help us identify important resistance mutations before they spread in clinics and agriculture.

Although our experiments were performed on the *C. albicans* gene, the conservation of the protein among orthologs may allow us to interpret the impact of mutations in other species. For instance, mutation F444L was recently identified and validated as contributing to fluconazole and voriconazole resistance in *C. auris*^67^. Here, we find that the homologous change F449L in *C. albicans* also confers resistance (*s* = 0.18 and 0.22, *P* < 0.0001). In addition, Williamson *et al.*^68^ tested the susceptibility of five *C. auris* variants V125A, F126L, Y132F, K143R and I466M (I471M in CaErg11) to fluconazole, voriconazole, itraconazole and posaconazole. We find similar resistance phenotypes for I471M and K143R (*s* = 0.11 to 0.43, *P* < 0.0001) but not for the other mutations. For instance, they^68^ found that V125A conferred a 2-fold decrease in susceptibility to voriconazole and itraconazole, but not to fluconazole and posaconazole while we classified this variant as resistant to all of the azoles we examined (*s* = 0.13 to 0.26, *P* < 0.0001). It is possible that standard methods used by the authors to quantify susceptibility are not sensitive enough to detect small variations, thereby explaining the observed differences. In the more distantly related mold *A. fumigatus,* G54E, V, R and W substitutions confer resistance to itraconazole and posaconazole while maintaining susceptibility to voriconazole^33^. The residue at position 54, orthologous to G65 in *C. albicans,* is located in the substrate entry channel. In accordance with Sagatova *et al.*^69^ who reproduced the mutations at the homologous position in *S. cerevisiae*, we did not detect an increase in azole resistance. We instead observed that two variants at position 54 have their growth impaired in control conditions and that all variants are deleterious in azoles. The substitution from glycine to larger amino acids could restrict entry channel access for drug and substrate. The disparities of phenotypes may arise due to differences in protein structure, with *Aspergillus* being a more distant species. Some slight changes could have occurred either on the branch leading to *Aspergillus* or the one leading to the yeast clade. Nonetheless, some resistance phenotypes are consistent between *C. albicans* and *A. fumigatus* mutations. For instance, *A. fumigatus* G138C and Y431C confer an approximately 10-fold increase in resistance to voriconazole, itraconazole and posaconazole^32^. Here, we also found that the homologous substitutions A149C and Y447C confer resistance to all of the azoles tested (*s* = 0.19 to 0.28, *P* < 0.0001). To help the community to perform such comparisons, we provide the selection coefficients and corresponding categories obtained for each CaErg11 variant at orthologous positions in fungal pathogens as a resource (Supplementary data 4).

In the future, it will be important to further investigate the different determinants of azole resistance mutations between *ERG11* orthologues and all azole drugs. For instance, azoles are also used in agriculture and evidence shows that azole-resistant *A. fumigatus* from the environment can infect patients leading to an increase in treatment failures^14,16,70^. This is alarming as it implies the presence of cross-resistance between agricultural and medical azoles^15,71,72^. The azoles used in agriculture are not the same as the ones used in clinics, but they have chemical similarities and they share the same mechanism of action. Given the extent of cross-resistance and the very limited tradeoff we measured here, the selective forces that represent the use of azoles in contexts such as in agriculture should be considered as a serious threat to the longevity of azoles for successfully treating some fungal infections^73,74^.

## Acknowledgement

We thank the members of the Landry lab for supportive advice and discussion on the project and Philippe Després and Romain Durand for sharing some bioinformatics material for data analysis. We also thank Aleeza Gerstein, Rong Shi and Sophie Gobeil for their advice and comments throughout the project.

## Author contributions

CRL, CB, IGA, LM and RSS designed research. CB, IGA, JB, SP, AKD, JS performed experiments. AP and CB generated in silico data. AF performed the population genomics analyses. CB analyzed data. CB produced figures. CB and CRL drafted the manuscript. All authors contributed to editing the manuscript.

## Financial support

This project was funded by a Canadian Institutes of Health Research (CIHR) Foundation grant number 387697 to CRL, a Fonds de recherche du Québec Nature et technologies (FRQNT) Team Grant (2022-PR-298169), a Genome Canada and Genome Quebec grant (6569 to CRL) and a Natural Sciences and Engineering Research Council of Canada (NSERC) Discovery Grant (RGPIN-2018-4914) to RSS. CRL holds the Canada Research Chair (Tier I) in Cellular Systems and Synthetic Biology. RSS holds the Canada Research Chair (Tier II) in Microbial Functional Genomics. CB was supported by fellowships from the Vanier Canada Graduate Scholarship agency, FRQNT, Fonds de Recherche du Québec Santé (FRQS), NSERC, EvoFunPath NSERC Collaborative Research and Training Experience (CREATE) program and Université Laval. JB was supported by fellowships from NSERC and FRQNT. LM was supported by a fellowship from the EvoFunPath NSERC CREATE program.

## Data availability

Raw sequencing reads from pooled competitions are available at BioProject PRJNA1044755. Read counts for DNA variants are available in Supplementary data 5, sample descriptions in Supplementary data 6 and overall read counts through quality control for every sample in Supplementary data 7. All files used in the analysis are available on GitHub at https://github.com/Landrylab/Bedard_et_al_2023.

## Code availability

All scripts for figures and analysis are available with their input and output files on GitHub at https://github.com/Landrylab/Bedard_et_al_2023.

## Methods

### Strains, antimicrobials and culture media

*Escherichia coli* MC1061 ([*araD139]_B/r_ Δ(araA-leu)7697 ΔlacX74 galK16 galE15(GalS) λ-e14-mcrA0 relA1 rpsL150(strR) spoT1 mcrB1 hsdR2*) was used for all cloning steps. *S. cerevisiae* R1158^75^ (*URA3*::CMV-tTA *MAT**a** his3-1 leu2-0 met15-0*) was the background strain used to construct *ScERG11*-DOX strain (Yeast strain construction section). *C. albicans* SC5314^76^ was the background strain used for genome editing and physiological profiling.

Nourseothricin (NAT, Cedarlane Labs : AB-102-25G), doxycycline (DOX, BioShop : DOX444.1), geneticin (G418, BioShop : GEN418.10) and ampicillin (AMP, BioShop : AMP201.25) were solubilized in water at a 1000X stock concentration, filtered and frozen at - 20 °C. Fluconazole (Cedarlane Labs : 11594-500), voriconazole (TCI America : V0116), itraconazole (Sigma-Aldrich : I6657), posaconazole (Sigma-Aldrich : 32103), isavuconazole (Sigma-Aldrich : SML2357), clotrimazole (TCI America : C28675G) were solubilized in dimethyl sulfoxide (DMSO) at 1000X stock concentration and stored at −20 °C.

For liquid cultures, bacteria were grown in LB medium (0.5% yeast extract, 1% tryptone and 1% sodium chloride) at 37 °C in a shaking incubator (250 rpm). For solid culture, bacteria were grown on 2YT medium with glucose and 2% agar (1% yeast extract, 1.6% tryptone, 0.5% sodium chloride and 0.2% glucose) and incubated at 37 °C. For bacterial transformations, medium was supplemented with 100 μg/mL AMP. Yeast were grown in YPD medium (1% yeast extract, 2% tryptone and 2% glucose) at 30 °C in a shaking incubator (250 rpm). On solid medium, 2% agar was added to YPD and yeasts were grown at 30 °C. Yeast transformants with pRS31N plasmids were selected on YPD medium supplemented with 100 μg/mL of NAT.

### Plasmids construction

All PCRs in this section were done using the KAPA HiFi HotStart polymerase (Roche : 07958897001). For all PCR amplification products used for cloning, 20 U of DpnI (New England Biolabs : R0176S) was added to the PCR products and incubated at 37 °C for 1 hour to remove the parental PCR DNA template. The digested amplicons were purified on magnetic beads (Axygen AxyPrep Mag PCR Clean-Up Kit). The vectors and inserts were assembled following the Gibson DNA assembly protocol^77^. Oligos and plasmids are listed in Supplementary data 8.

*Candida albicans ERG11* (*CaERG11*) was expressed from a yeast centromeric plasmid pRS31N-*CaERG11* constructed as follows. First, we created a plasmid containing the *S. cerevisiae ERG11* (*ScERG11*) sequence, including its promoter and terminator. We assembled pRS31N-*ScERG11* as follows. The plasmid pRS31N^78^ containing AMP and NAT resistance markers was amplified using this PCR mix and cycles:

|  | **Name** | **Concentration** | **Volume (μL)** |
| --- | --- | --- | --- |
| DNA | pRS31N | 10 ng/μL | 1.00 |
| Buffer | Fidelity buffer | 5X | 5.00 |
| dNTPs |  | 10 mM | 0.75 |
| Oligo F | OP252-A10 | 10 μM | 0.75 |
| Oligo R | OP252-B10 | 10 μM | 0.75 |
| Enzyme | KAPA | 1 U/μL | 0.30 |
| Water |  |  | 16.45 |
|  |  | **Total volume** | 25.00 |

| **Time** | **Temperature** | **Number of cycles** |
| --- | --- | --- |
| 5 min | 95°C | 1x |
| 20 sec | 98°C | 35x |
| 15 sec | 66°C |  |
| 3 min | 72°C |  |
| 3 min | 72°C | 1x |
| Hold | 10°C |  |

The *S. cerevisiae ERG11* native promoter (878 base pairs), coding sequence (1593 base pairs) and terminator (319 base pairs) were amplified from the pMoBY-*ERG11* plasmid^79^ with primers adding homology to the pRS31N plasmid cloning site:

|  | **Name** | **Concentration** | **Volume (μL)** |
| --- | --- | --- | --- |
| DNA | pMoBY-*ERG11* | 10 ng/μL | 1.00 |
| Buffer | Fidelity buffer | 5X | 5.00 |
| dNTPs |  | 10 mM | 0.75 |
| Oligo F | OP252-C10 | 10 μM | 0.75 |
| Oligo R | OP252-D10 | 10 μM | 0.75 |
| Enzyme | KAPA | 1 U/μL | 0.30 |
| Water |  |  | 16.45 |
|  |  | **Total volume** | 25.00 |

| Time | Temperature | Number of cycles |
| --- | --- | --- |
| 5 min | 95°C | 1x |
| 20 sec | 98°C | 5x |
| 15 sec | 61°C |  |
| 2 min | 72°C |  |
| 20 sec | 98°C | 30x |
| 15 sec | 68°C |  |
| 2 min | 72°C |  |
| 3 min | 72°C | 1x |
| Hold | 10°C |  |

The vector and the insert were assembled to obtain pRS31N-*ScERG11*.

For the second step to construct pRS31N-*CaERG11*, pRS31N-*ScERG11* was amplified to remove the *ScERG11* sequence and keep the native promoter and terminator:

|  | Name | Concentration | Volume (µL) |
| --- | --- | --- | --- |
| DNA | pRS31N-*ScERG11* | 10 ng/µL | 1.00 |
| Buffer | Fidelity buffer | 5X | 5.00 |
| dNTPs |  | 10 mM | 0.75 |
| Oligo F | OP252-G8 | 10 µM | 0.75 |
| Oligo R | OP252-H8 | 10 µM | 0.75 |
| Enzyme | KAPA | 1 U/µL | 0.30 |
| Water |  |  | 16.45 |
|  |  | **Total volume** | 25.00 |

| Time | Temperature | Number of cycles |
| --- | --- | --- |
| 5 min | 95°C | 1x |
| 20 sec | 98°C | 35x |
| 15 sec | 63°C |  |
| 6 min | 72°C |  |
| 8 min | 72°C | 1x |
| Hold | 10°C |  |

To ensure the expression of *CaERG11* in *S. cerevisiae* (*C. albicans* decodes the CTG codon differently^80^), the *CaERG11* sequence (from SC5314 strain^76^, Candida Genome Database (CGD): C5_00660C_A) was codon optimized for *S. cerevisiae* (GenScript). The sequence was synthetized by GenScript and amplified by adding homology to the *ScERG11* promoter and terminator, so it could replace the *ScERG11* coding sequence in pRS31N-*ScERG11*:

|  | **Name** | **Concentration** | **Volume (μL)** |
| --- | --- | --- | --- |
| DNA | pUC19-*CaERG11* | 10 ng/μL | 1.00 |
| Buffer | Fidelity buffer | 5X | 5.00 |
| dNTPs |  | 10 mM | 0.75 |
| Oligo F | OP252-C8 | 10 μM | 0.75 |
| Oligo R | OP252-D8 | 10 μM | 0.75 |
| Enzyme | KAPA | 1 U/μL | 0.30 |
| Water |  |  | 16.45 |
|  |  | **Total volume** | 25.00 |

| **Time** | **Temperature** | **Number of cycles** |
| --- | --- | --- |
| 5 min | 95°C | 1x |
| 20 sec | 98°C | 5x |
| 15 sec | 60°C |  |
| 1 min | 72°C |  |
| 20 sec | 98°C | 30x |
| 15 sec | 68°C |  |
| 1 min | 72°C |  |
| 2 min | 72°C | 1x |
| Hold | 10°C |  |

The vector and the insert were cloned to obtain pRS31N-*CaERG11*.

The variant library (see below for library design and construction) was cloned in pRS31N-Barcode, a plasmid containing a NdeI restriction site, a 20N barcode and universal priming sites for subsequent amplifications. The barcodes were not used in this project, but were added in anticipation of future experimentation. To construct this plasmid, pRS31N-*ScERG11* was amplified without *ScERG11* and the terminator. The forward oligo added the priming sites and the barcode:

|  | **Name** | **Concentration** | **Volume (μL)** |
| --- | --- | --- | --- |
| DNA | pRS31N-*ScERG11* | 10 ng/μL | 5.00 |
| Buffer | Fidelity buffer | 5X | 5.00 |
| dNTPs |  | 10 mM | 0.75 |
| Oligo F | CLOP259-F3 | 10 μM | 2.00 |
| Oligo R | CLOP252-H8 | 10 μM | 2.00 |
| Enzyme | KAPA | 1 U/μL | 0.30 |
| Water |  |  | 9.95 |
|  |  | **Total volume** | 25.00 |

| Time | Temperature | Number of cycles |
| --- | --- | --- |
| 5 min | 95°C | 1x |
| 20 sec | 98°C | 18x |
| 15 sec | 62°C |  |
| 5 min | 72°C |  |
| 3 min | 72°C | 1x |
| Hold | 10°C |  |

From pRS31N-*ScERG11*, the terminator was amplified with oligos adding a NdeI restriction site and homology to the vector:

|  | Name | Concentration | Volume (μL) |
| --- | --- | --- | --- |
| DNA | pRS31N-*ScERG11* | 10 ng/μL | 1.00 |
| Buffer | Fidelity buffer | 5X | 5.00 |
| dNTPs |  | 10 mM | 0.75 |
| Oligo F | CLOP259-D3 | 10 μM | 0.75 |
| Oligo R | CLOP259-E3 | 10 μM | 0.75 |
| Enzyme | KAPA | 1 U/μL | 0.30 |
| Water |  |  | 16.45 |
|  |  | **Total volume** | 25.00 |

| Time | Temperature | Number of cycles |
| --- | --- | --- |
| 5 min | 95°C | 1x |
| 20 sec | 98°C | 5x |
| 15 sec | 64°C |  |
| 15 sec | 72°C |  |
| 20 sec | 98°C | 30x |
| 15 sec | 70°C |  |
| 15 sec | 72°C |  |
| 3 min | 72°C | 1x |
| Hold | 10°C |  |

The vector and the insert were cloned to obtain pRS31N-Barcode. To preserve barcode diversity, we added 5 mL of 2YT medium to the plates with colonies following Gibson DNA assembly, scraped them with a glass rake and used 1.5 mL of this mix to extract plasmids.

### Yeast strain construction

The construction of the *ERG11* repressible strain of *S. cerevisiae* (*ScERG11*-DOX) was based on the Yeast Tet-Promoters Hughes collection (yTHC)^39^. The endogenous promoter of *ERG11* (50 bp before the ATG codon of ERG11, chrVIII:121683..121733) was replaced with a TET-titratable promoter repressible with doxycycline (DOX). The *ScERG11*-DOX strain was constructed as follows. The *KANMX*-tetO_7_ cassette was amplified from pKB33^81^:

|  | Name | Concentration | Volume (μL) |
| --- | --- | --- | --- |
| DNA | pKB33 | 10 ng/μL | 0.75 |
| Buffer | Reaction buffer | 5X | 5.00 |
| dNTPs |  | 10 mM | 0.50 |
| Oligo F | CLOP258-G6 | 10 μM | 1.25 |
| Oligo R | CLOP258-D7 | 10 μM | 1.25 |
| Enzyme | Q5* | 1 U/μL | 0.25 |
| Water |  |  | 16.00 |
|  |  | **Total volume** | 25.00 |

\*Q5 High-Fidelity DNA Polymerase (NEB : M0491L)

| Time | Temperature | Number of cycles |
| --- | --- | --- |
| 5 min | 95°C | 1x |
| 20 sec | 98°C | 5x |
| 15 sec | 66°C |  |
| 2 min | 72°C |  |
| 20 sec | 98°C | 30x |
| 15 sec | 70°C |  |
| 2 min | 72°C |  |
| 3 min | 72°C | 1x |
| Hold | 10°C |  |

Competent cells of *S. cerevisiae* R1158^75^ were transformed with the amplified *KANMX*-tetO_7_ cassette using a standard lithium acetate transformation protocol^82^. G418 was added to the medium at a concentration of 200 μg/mL. The resulting *ScERG11*-DOX strain has the genotype pERG11::kanR-tetO_7_-TATA *URA3*::CMV-tTA *MAT**a** his3-1 leu2-0 met15-0*.

### Dose-response curves

Yeasts were grown in 96-well plates containing 200 μL of YPD medium with NAT, DOX and azole at a starting optical density at 600 nm (OD_600_) of 0.01. OD_600_ was measured in a Tecan Infinite M Nano (Tecan Life Sciences) every 15 minutes for 24 hours without agitation and at 30 °C. The maximum growth rates were calculated from growth curves across a minimum of 17 drug concentrations, in triplicate for fluconazole and in duplicate for other azoles. The maximum growth rates were transformed into relative inhibition to get dose-response curves. Hill equation was fitted to the dose-response curves to determine the minimal inhibitory concentration at 50% (IC_50_). The analysis was based on Durand *et al*^83^.

### Library design

The selection of positions to mutate to create the CaErg11 deep mutational scanning library was done using the ScErg11 structures. The crystal structures of ScErg11 bound to lanosterol (4LXJ), fluconazole (4WMZ) and itraconazole (5EQB) were taken from the Protein Data Bank (PDB) and used to calculate the distances between the residues and the substrates and the heme molecule. The distances were compared with residues with known or putative resistance mutations from the MARDy database^84^. A radius of 12 Å covering the active site and most residues with known resistance mutations was selected. An alignment between ScErg11 and CaErg11 using the Biopython toolkit (pairwise, global, BLOSUM62)^85^ was used to select the homologous residues in the *C. albicans* ortholog with a custom script. ScErg11 and CaErg11 sequences have a sequence identity of 62.7%. Consequently, their 3D structures are very similar (Extended Data Fig. 2). The identity between ScErg11 and CaErg11 for the 206 residues in the 12 Å radius is 85.9%. The selected residues are listed in the Supplementary data 1.

In the CaErg11 variant library, each of the 206 targeted residues were replaced by the 19 non-wild type amino acids encoded by two codons, when possible (methionine and tryptophan are encoded by only one codon, Supplementary data 2). For each amino acid, we selected the two most frequently used codons in the *ScERG11* ortholog to minimize the potential impact of codon usage on expression. As controls, a stop codon was introduced at every 20 positions along the sequence. A total of 7,839 DNA variants were designed. We ordered the synthesized DNA variants from Twist Bioscience (QC report, Supplementary data 9). Positions 92, 228, 232, 233 and 234 failed to be synthesized, so they are missing from the library. The library used covers a total of 3,830 amino acid variants. The library was split into four fragments (F1, F2, F3 and F4) of around 400 base pairs with overlapping positions. Fragment 1 goes from residue 44 to 176, fragment 2 goes from residue 147 to 288, fragment 3 goes from residue 269 to 413 and fragment 4 goes from residue 386 to 520.

### Library construction

The variant sequences have a flanking region of 35 base pairs from the native *ScERG11* promoter to clone into pRS31N-Barcode. To limit the risk of losing variant diversity, we clone the variants position-by-position. The variant sequences were amplified with a reverse primer adding homology to the *ScERG11* terminator using the following PCR mix and conditions:

|  | Name | Concentration | Volume (µL) |
| --- | --- | --- | --- |
| DNA | Twist DNA variants | 1 ng/µL | 1.00 |
| Buffer | Fidelity buffer | 5X | 5.00 |
| dNTPs |  | 10 mM | 0.75 |
| Oligo F | CLOP259-D3 | 10 µM | 0.75 |
| Oligo R | CLOP259-E3 | 10 µM | 0.75 |
| Enzyme | KAPA | 1 U/µL | 0.30 |
| Water |  |  | 16.45 |
|  |  | **Total volume** | 25.00 |

| **Time** | **Temperature** | **Number of cycles** |
| --- | --- | --- |
| 5 min | 95°C | 1x |
| 20 sec | 98°C | 5x |
| 15 sec | 60°C |  |
| 1 min | 72°C |  |
| 20 sec | 98°C | 13x |
| 15 sec | 68°C |  |
| 1 min | 72°C |  |
| 2 min | 72°C | 1x |
| Hold | 10°C |  |

pRS31N-Barcode was digested with NdeI (NEB : R0111S) for 2 hours at 37°C and inactivated for 20 minutes at 65°C:

| **Component** | **Concentration** | **Volume (µL)** |
| --- | --- | --- |
| pRS31N-Barcode | 93.1 ng/µL | 21.5 |
| CutSmart™ Buffer | 10X | 4.0 |
| NdeI | 20 U/µL | 2.0 |
| Water |  | 12.5 |
|  |  | 40.0 |

Variant sequences were cloned into the NdeI digested pRS31N-barcode plasmid by Gibson DNA assembly^77^. We added 5 mL of 2YT medium to the plates, they were scraped with a glass rake and 1.5 mL was used to extract plasmids for yeast transformations.

The plasmid variant library was transformed position-by-position in competent *ScERG11*-DOX yeast cells using a standard lithium acetate transformation protocol^82^. We added 5 mL of YPD medium to the plates and scraped with a glass rake. All CaErg11 library positions were finally pooled by fragment in equivalent of OD_600_ units and archived in 25% glycerol at −80°C (Extended Data Fig. 3).

### Library quality control

The yeast library was grown by pooling fragments and growing in liquid YPD+NAT medium overnight. We aliquoted 5 OD_600_ units of cells into 1.5 mL microtubes and spun down at 600 x g for 2 minutes. Then, after supernatant removal, 1 mL of YPD was added to resuspend the pellets and cells were spun down again at 600 x g for 2 minutes. The supernatant was removed by aspiration and the resulting pellets were kept at −80°C until they were processed for plasmid extraction.

Plasmids were retrieved from yeast using the Zymoprep Yeast Plasmid Miniprep II kit (Zymo Research : D2004). The protocol was modified according to Starr et al., 2020^86^ and from our own tests to increase yield. To frozen pellets, 200 μL of solution 1 was added. Then, 30 U of Zymolyase™ (from the kit and from BioShop : ZYM002) was added. Cells were resuspended by light vortexing and incubated at 37 °C for 2 hours. At the end of the incubation period, tubes were lightly vortexed and frozen for at least 20 minutes at −80°C, and up to overnight. Frozen tubes were thawed at 37°C for 3 minutes. After, 200 μL of solution 2 was added and mixed. Then, 400 μL of solution 3 was added, mixed and tubes were centrifuged at maximum speed for 3 minutes. The supernatant was transferred to the Zymo-Spin-I column (around 800 μL) and spun for 30 seconds at maximum speed. To the column in a collection tube, 550 μL of wash buffer with ethanol was added. Column was centrifuged for 1 minute 30 seconds at maximum speed. The column was then placed into a new 1.5 mL microfuge tube, 10 μL of Tris 10 mM pH 8 was added, and the tube was spun for 1 minute at maximum speed. The resulting elution was conserved at −20°C.

Starting with 2 μL of yeast miniprep, *CaERG11* was amplified with a different primer pair for each of the four fragments (F1: CLOP279-A6 + CLOP279-B8, F2: CLOP279-C6 + CLOP279-D8, F3: CLOP279-E6 + CLOP279-F8, F4: CLOP279-G6 + CLOP279-H8) and annealing temperature (**A** = F1: 61°C, F2 :60°C, F3: 58°C, F4: 62°C):

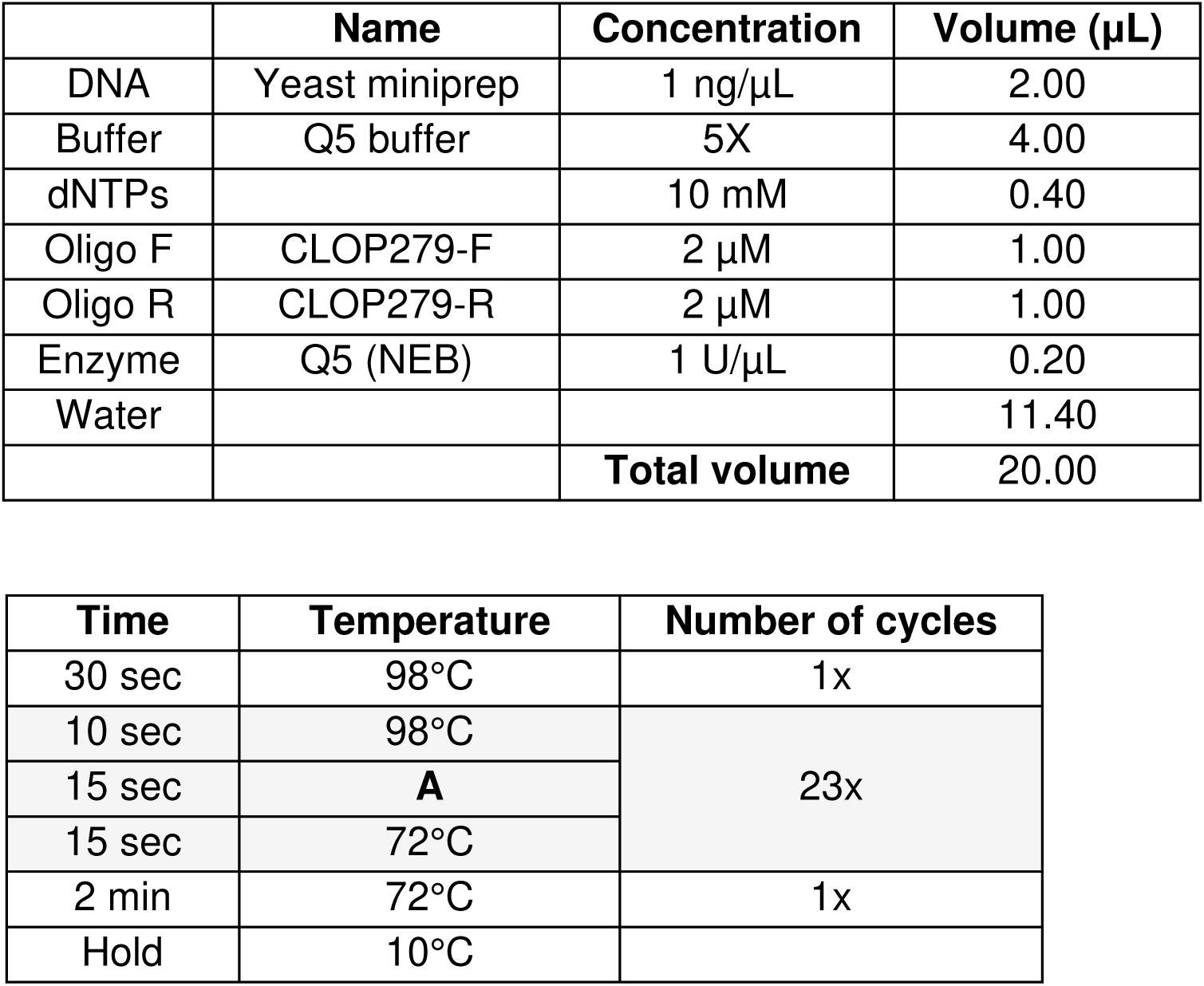

The amplicons were diluted 1:2,500 and used as a template for a second PCR adding Illumina sequencing adaptors. The reverse primer was unique for each fragment (F1: CLOP225-A1, F2: CLOP225-B1, F3: CLOP225-C1, F4: CLOP225-D1):

|  | Name | Concentration | Volume (µL) |
| --- | --- | --- | --- |
| DNA | *CaERG11* fragments | 1/2500 | 1.00 |
| Buffer | Fidelity buffer | 5X | 4.00 |
| dNTPs |  | 10 mM | 0.60 |
| Oligo F | CLOP224-A1 | 2 µM | 3.00 |
| Oligo R | CLOP225-R | 2 µM | 3.00 |
| Enzyme | Kapa | 1 U/µL | 0.13 |
| Water |  |  | 8.27 |
|  |  | **Total volume** | 20.00 |

| Time | Temperature | Number of cycles |
| --- | --- | --- |
| 3 min | 98°C | 1x |
| 30 sec | 98°C | 18x |
| 15 sec | 61°C |  |
| 35 sec | 72°C |  |
| 1 min | 72°C | 1x |
| Hold | 10°C |  |

Amplicons were purified on magnetic beads (Axygen AxyPrep Mag PCR Clean-Up Kit).

High-throughput sequencing was performed at the Centre de recherche du CHU de Québec - Université Laval sequencing platform (CHUL, Québec) on an Illumina NovaSeq 6000 in 2×250 bp paired-end reads. The library was sequenced at roughly 100 reads per variant (F1: 7 million, F2: 3 million, F3: 5 million, F4: 5 million) to estimate diversity and confirm the presence of every variant. All mutations were confirmed to be present in the library, but a handful appeared to be present at very low copy number (109CAG: 24 reads, 374ACT: 66 reads, 374CAA: 46 reads, 374GTC: 51 reads and 472TGT: 86 reads).

### Pooled competition assay and high-throughput sequencing

We competed the *CaERG11* variants with and without azoles in a medium containing doxycycline (10 μg/mL) to repress the endogenous copy of Sc*ERG11* and NAT (100 μg/mL) for plasmid maintenance. Since antifungals were solubilized in DMSO, the control condition also contained 0.1% DMSO. Azoles were used at concentrations inhibiting around 50% of the wild-type growth (Extended Data Fig. 5) (fluconazole: 10.5 µg/mL, voriconazole: 0.12 µg/mL, itraconazole: 0.10 µg/mL, posaconazole: 0.10 µg/mL, isavuconazole: 0.10 µg/mL and clotrimazole: 0.30 µg/mL). We had to use lower concentrations for itraconazole and isavuconazole than anticipated by our IC_50_ calculations as growth did not resume in these conditions upon dilutions in the second round of competition.

Each of the four pools of the library were grown overnight in YPD+NAT starting with around 16,000 cells per variant and 2% of wild-type sequence (Timepoint 0, TP0). The saturated cultures were transferred to fresh YPD+NAT+DOX+azole at a starting OD_600_ of 0.05 and grew until an OD_600_ of 0.80 was reached (Timepoint 1, TP1). We did another transfer in fresh YPD+NAT+DOX+azole at starting OD_600_ of 0.05, and let them grow until they reached an OD_600_ of 0.80 (Timepoint 2, TP2) to achieve eight generations of competition (Extended Data Fig. 4). For the no antifungal condition, the same protocol was used, but we added a third timepoint for a total of 12 generations.

Aliquoted pools of TP0 were prepared using the same protocol as in the “Library quality control” section. For TP2 (TP3 for the control condition), we aliquoted 5 OD_600_ of cells in 15 mL tubes and centrifuged at 916 x g for 5 min. Then, after supernatant removal, 1 mL of YPD was added to resuspend the pellets. The resuspensions were transferred in a 1.5 mL microtube and cells were spun down again at 600 x g for 2 min. The supernatant was removed by aspiration and the resulting pellets were kept at −80°C until they were processed for plasmid extraction. Yeast minipreps and sequencing were done using the same protocol as in the “Library quality control” section. We used the primers listed in the sample files (Supplementary data 6) and sequenced at roughly 100 reads per variant (5 million reads per fragment, except for the control condition and in fluconazole, F1= 7 million and F2= 3 million). Only variants from the TP2 are sequenced since after only four generations (TP1), the distribution of variant frequency changes is narrow, limiting the correct association of the genotypes with the phenotypes.

### High-throughput sequencing analysis and variants count

High-throughput sequencing results were analyzed using custom scripts based on Després *et al*.^21^ Each fragment has a different script but following the same steps. In the first part, quality control was done on demultiplexed reads with FastQC^87^, reads were merged using PANDAseq^88^ and trimmed according to oligos length and aggregated with VSEARCH^89^. Reads were then aligned to their corresponding *CaERG11* fragment using Needle from EMBOSS^90^. In the second part, Needle output files obtained were parsed to find mutations and count variants. As the library contains only single amino acid variants, we considered only reads where all the mutations detected were in the same codon when more than one mutation was detected.

### Selection coefficient and characterization of the variants

The raw variant read counts were normalized by the total number of reads in their sequencing pool. The normalized read counts (N) of the initial timepoint (TP0) and the final timepoint (TP2) for the missense variants (mut) and the median normalized read counts for the variants with wild-type synonymous codons (med_WT) were used with the number of generations (G) to estimate a selection coefficient (*s*) of the variants using this equation^91^:

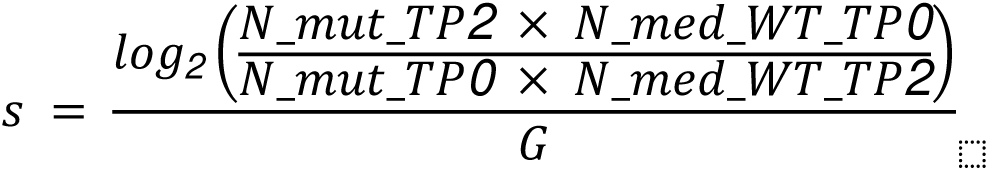

We examined correlation between replicates (Extended data Fig.6), fragment overlaps (Extended data Fig.7) and synonymous codons (Extended data Fig.8) to measure the reproducibility of the results. Only F4 of the fluconazole condition was not done at the same moment as the others, hence it was not as linearly correlated with F3 as we saw for other comparisons. This could have been caused by slight differences between batches of experiments. To correct for this effect, we built an interpolation model to get new F4 values following a linear regression with F3. The sequencing failed for 2 samples and they had to be removed (Replicate A of F1 for voriconazole and Replicate C of F4 for itraconazole).

We classified the variants into three classes: beneficial, wild-type-like and deleterious. To do so, we compared the variants with wild-type synonymous codons to missense variants with two-sided independent *t-test* and corrected for multiple comparisons with the Benjamini-Hochberg method. Variants with a false discovery rate (FDR) corrected *p-value* lower than 0.01 and with a positive selection coefficient were classified as beneficial, and those with a negative selection coefficient, as deleterious. All variants with a p-value higher than 0.01 were classified as neutral.

### Validations by site-directed mutagenesis

The validation variants (Supplementary data 8) were constructed by site-directed mutagenesis on the pRS31N-*CaERG11* plasmid. The mutagenesis primers designed were 40 base pair long and identical to the sequence of the plasmid except for the bases we mutated:

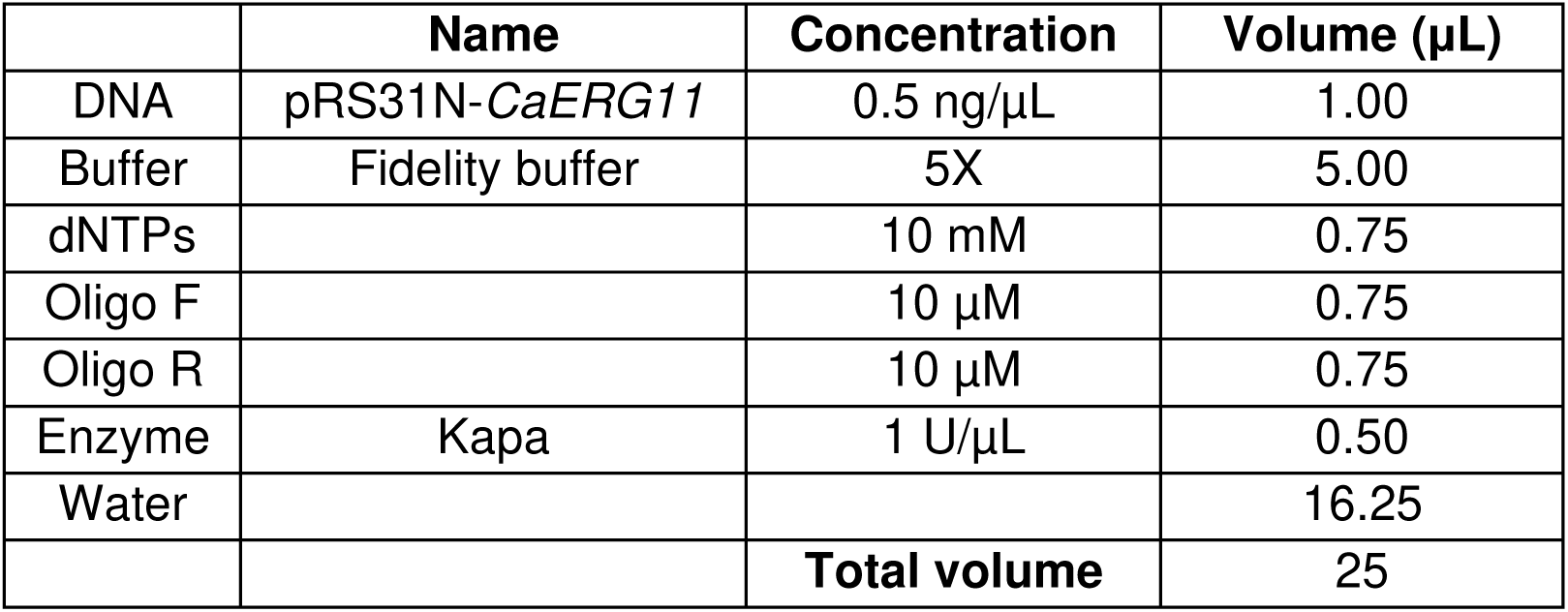
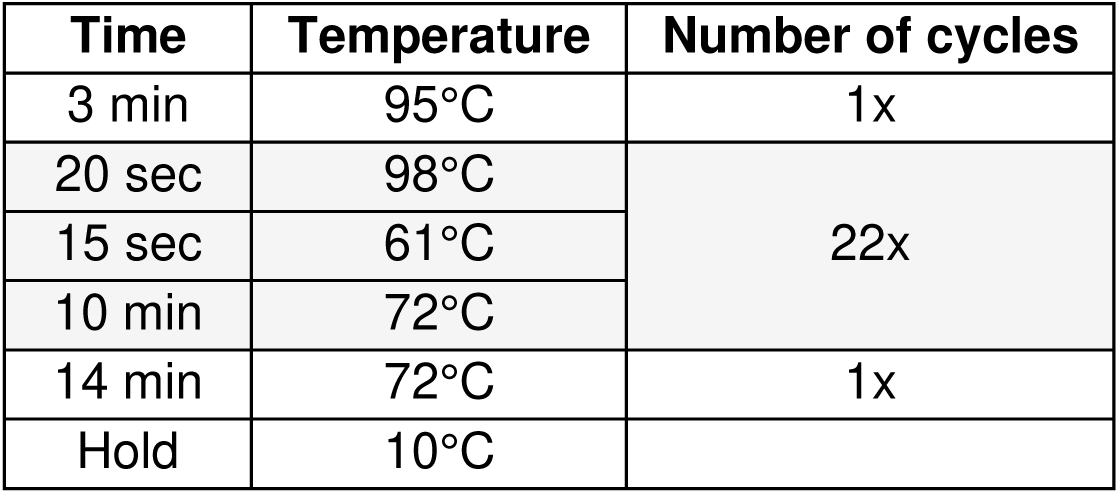

Every product of amplification was digested with 20 U of DpnI and incubated at 37°C for 1 hour. Transformation in competent bacteria was done to retrieve the modified plasmids and the mutations were confirmed by Sanger sequencing. Plasmids were transformed into competent *ScERG11*-DOX cells using a standard lithium acetate transformation protocol^82^.

Yeast were grown in YPD+DOX+NAT+0.1% DMSO and azoles as described in the dose-response curves section. Azole concentrations corresponding to the IC_50_ were used. The area under the curve (AUC) was extracted from the resulting growth curves using the composite trapezoidal rule.

### Quantification of Erg11 abundance

We chose a subset of fluconazole resistant validation variants along with susceptible controls to quantify Erg11 abundance. We cloned the mEGFP downstream of *CaERG11* coding sequence in the pRS31N-*CaERG11* plasmids (wild-type and mutated). pRS31N-*CaERG11* was amplified with primers removing *CaERG11* stop codon as followed:

|  | Name | Concentration | Volume (μL) |
| --- | --- | --- | --- |
| DNA | pRS31N-*CaERG11* | 10 ng/μL | 1.00 |
| Buffer | Fidelity buffer | 5X | 5.00 |
| dNTPs |  | 10 mM | 0.75 |
| Oligo F | CLOP321-F2 | 10 μM | 0.75 |
| Oligo R | CLOP321-G2 | 10 μM | 0.75 |
| Enzyme | Kapa | 1 U/μL | 0.50 |
| Water |  |  | 16.25 |
|  |  | **Total volume** | 25 |

| Time | Temperature | Number of cycles |
| --- | --- | --- |
| 5 min | 95°C | 1x |
| 20 sec | 98°C | 35x |
| 15 sec | 63°C |  |
| 8 min | 72°C |  |
| 10 min | 72°C | 1x |
| Hold | 10°C |  |

The mEGFP was amplified from F65V-34a-mEGFP plasmid as followed using primer adding homology to *CaERG11* and to the native *ScERG11* terminator:

|  | Name | Concentration | Volume (μL) |
| --- | --- | --- | --- |
| DNA | F65V-34a-mEGFP | 10 ng/μL | 1.00 |
| Buffer | Fidelity buffer | 5X | 5.00 |
| dNTPs |  | 10 mM | 0.75 |
| Oligo F | CLOP321-D2 | 10 μM | 0.75 |
| Oligo R | CLOP321-E2 | 10 μM | 0.75 |
| Enzyme | Kapa | 1 U/μL | 0.50 |
| Water |  |  | 16.25 |
|  |  | **Total volume** | 25 |

| Time | Temperature | Number of cycles |
| --- | --- | --- |
| 5 min | 95°C | 1x |
| 20 sec | 98°C | 5x |
| 15 sec | 60°C |  |
| 45 sec | 72°C |  |
| 20 sec | 98°C | 30x |
| 15 sec | 68°C |  |
| 45 sec | 72°C |  |
| 2 min | 72°C | 1x |
| Hold | 10°C |  |

Each amplicon was digested with 20 U of DpnI and incubated at 37°C for 1 hour. The digested amplicons were purified on magnetic beads and vectors and inserts were assembled following the Gibson DNA assembly protocol^77^. Plasmids were transformed into competent *ScERG11*-DOX cells using a standard lithium acetate transformation protocol^82^.

Yeasts were grown overnight in a 96 deep-well plate with shaking at 30°C in SC[MSG]-tryptophan+NAT medium (synthetic complete medium: 0.17 % yeast nitrogen base without amino acids and ammonium sulfate, 0.1 % monosodium glutamate, 2 % glucose and amino acids drop-out without tryptophan). The saturated cultures were diluted to an OD_600_ of 0.15 in the same medium and incubated again at 30°C with agitation until an OD_600_ of 0.5 was reached. Then, cells were diluted to an approximate concentration of 500 cells/µL in sterile ddH_2_O. The GFP fluorescence was measured with a Guava flow Cytometer (Millipore, USA). We normalized the GFP fluorescence signal with cell size using the FSC-H value and we filtered to conserve cells with a GFP signal above the median plus 2.5 standard deviations of the control without GFP.

### Candida albicans genome editing

We constructed pRS118, a plasmid derived from a previously described Candida-optimized Cas9 plasmid (pRS252)^92,93^. An sgRNA cloning locus (containing sgRNA promoter, SNR52 and the sgRNA tail) was cloned into pRS252 to generate pRS118. To create the H468T mutation in the *ERG11* gene, CRISPR-Cas9 was first used to mediate a double-stranded break and then the cells were supplied with the desired repair template. The sgRNA (oRS1335) for CaErg11 is a 20 bp sequence that was designed using the sgRNA design tool Eukaryotic Pathogen CRISPR gRNA Design Tool (EuPaGDT; http://grna.ctegd.uga.edu) which suggests guides based on specificity and efficiency^94^. This sgRNA targeted 378 bp downstream of the *ERG11* gene and was ordered from Integrated DNA Technologies (IDT). The sgRNA was then duplexed and cloned in the *C. albicans*-specific CRISPR-Cas9 plasmid, pRS118, using SapI-specific Golden Gate cloning, as described in Wensing *et al*.^95^ to generate pRS684. The repair template (gRS92) was ordered as a 971 bp gBlock gene fragment from IDT. This DNA fragment contains the mutations (CAT468ACC) and the modified PAM sequence (AGG to CGA) that prevents further cutting by pRS118’s Cas9 upon successful insertion of gRS92.

The repair template was amplified by PCR using the following conditions:

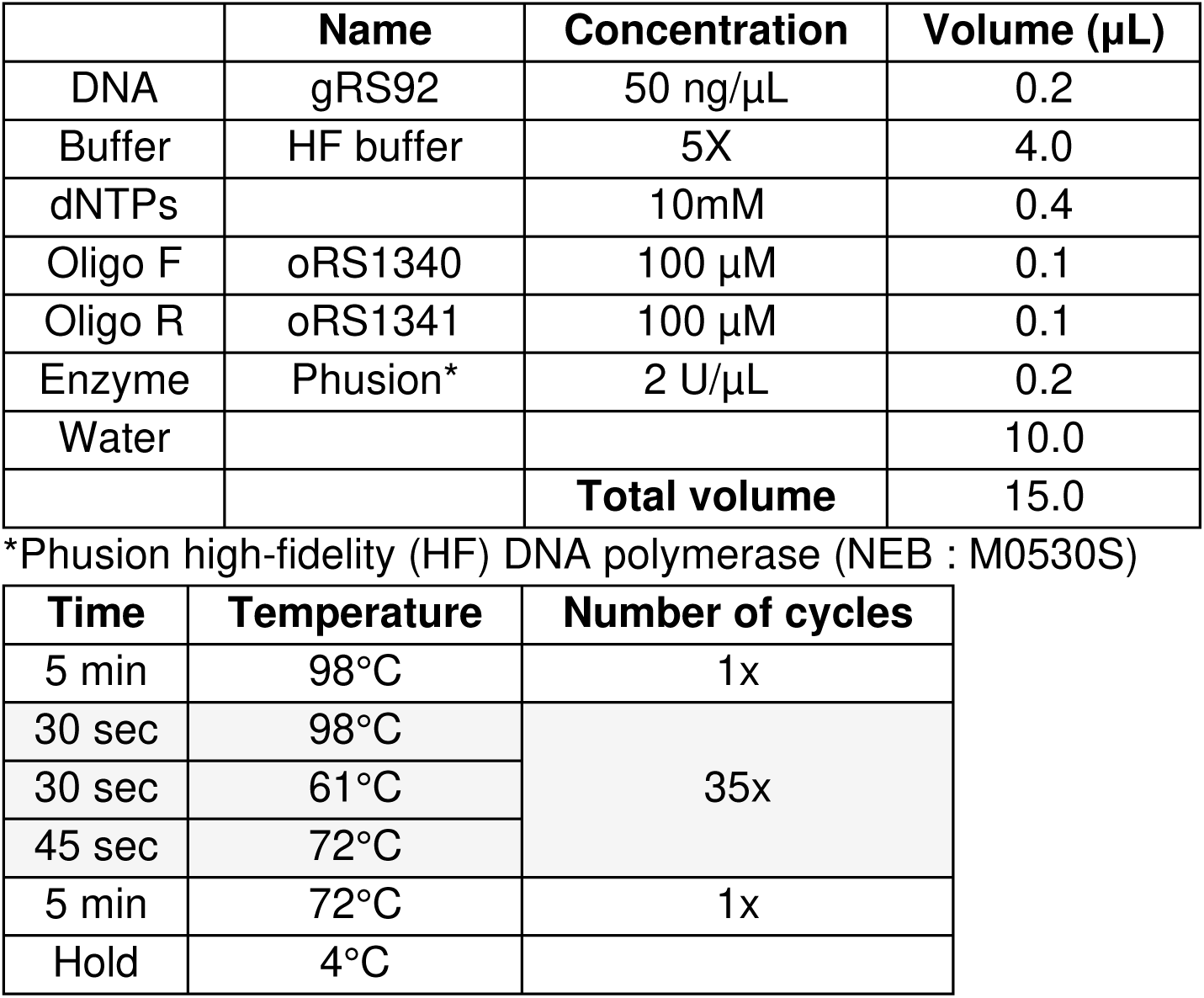

A lithium acetate transformation protocol was used to transform pRS684 into the SC5314 strain^95^. PacI-linearized plasmid and 100 µl of the amplified repair template was added to SC5314 in the LiAc transformation mix, heat shocked and recovered on YPD medium plates supplemented with 250 μg/mL of NAT^95^. Successful transformants with the plasmid inserted into the *NEUT5L* locus were confirmed using PCR. Sanger sequencing was used to confirm the *ERG11* heterozygous genotype for H468T of three colonies.

We confirmed the resistance phenotype of the edited strains (three H468H/T mutants and two wild-type controls) by spot assays. Cell density was adjusted at 1 of OD_600_ and diluted 1/5 four times. We spotted 5 μL on YPD media (YPD + DMSO, YPD + 2 μg/mL fluconazole and YPD + 4 μg/mL fluconazole) and incubated at 30°C for 24h.

### *Candida albicans* experimental evolution

A total of 12 CRISPR-Cas9 H468T *ERG11* genome edited *C. albicans* populations (four replicates per strain) were evolved for four days (around 35 generations) in the presence of increasing fluconazole concentrations. Cells were first grown overnight in a 96-well plate at 30°C in YPD medium (TP0). In the morning, yeasts were diluted 1/400 in 200 μl of fresh YPD+1μg/mL fluconazole medium (TP1) and grown at 30°C for 24h. The same manipulations were done for three more days, but with 1.5, 2 and 4 μg/mL of fluconazole (TP2, TP3 and TP4). Cells from each well were archived at −80°C in 50% glycerol every morning and cells from TP0 and TP4 were streaked on solid YPD medium to get individual colonies. One random colony for six of the TP0 and TP4 evolved populations (two populations per strain) was sent to Sanger sequencing to assess zygosity at the *ERG11* H468 locus.

Growth in the absence and in the presence of fluconazole (4 μg/mL) was measured for wild-type strains, for every H468H/T heterozygous strains sequenced from TP0 and for every H468T homozygous strains sequenced from TP4. Yeasts were incubated at 30°C in 96-well plates containing 200 μL of YPD medium at a starting optical OD_600_ of 0.05. OD_600_ was measured in a BioTek Synergy HTX Multimode Reader (Agilent) every 15 minutes for 24 hours without agitation.

### In silico data and bioinformatic analysis

We ran GEMME^48^ directly on the webserver (<u>GEMME (upmc.fr)</u>). As suggested in the instructions, we generated the multiple sequence alignment using MMseqs2 via ColabFold (A3M format)^96^ with CaErg11 sequence (CGD : C5_00660C_A) and uploaded the A3M file. We kept all default parameters (Number of iterations : 2 and Maximum number of sequences : 20,000). We used the data from the combined model (normPred_evolCombi.txt).

We used the MutateX package^97^ to run an in silico mutation scan of CaErg11 with foldX 5^49,98^. We used CaErg11 structure bound to itraconazole (PDB ID : 5V5Z) for this analysis. Heme iron and itraconazole were parameterized with the pyFoldX paramx package^99^ generating JSON files for those small molecules to be recognized by foldX. MutateX protocol runs RepairPDB, BuildModel and AnalyseComplexe commands from foldX to compute changes in protein free energy. The number of runs was set to 5. We analyzed interaction energy between the protein and antifungal drug as well as with the heme cofactor. Foldx ddG values higher than 5 were rounded to 5. Positions 432, 434 and 436 are absent from the crystalized protein structure, therefore, it was impossible to get ΔΔG values for them.

Whole genome sequence Illumina reads of *C. albicans* were downloaded from NCBI for four studies: Hirakawa et al. 2015^50^ (21 genomes), Chew et al.^51^ 2021 (14 genomes), Szarvas et al. 2021^53^ (30 genomes), and Gong et al. 2023^52^ (370 genomes). Reads were checked for quality with fastqc v0.12^100^, trimmed with Trimmomatic v0.39^101^, and mapped to the A haplotype of *C. albicans* SC5314 genome assembly A22-s07-m01-r183 (Candida Genome Database, accessed in August 2023) using bwa-mem2 v2.2.1^102^. Reads were realigned around INDELs with samtools calmd v1.17^103^ with options “-bAQr” and read duplicates were marked with picard’s MarkDuplicates v3.0.0. Details of the mapping pipeline are in https://github.com/aniafijarczyk/read-map-call. Single nucleotide polymorphisms were called using bcftools v1.17 with mpileup options: “mpileup -C50 -min-MQ 4 -min-BQ 13 --skip-any-set 1796 -a FORMAT/AD,FORMAT/ADR,FORMAT/ADF,FORMAT/DP” and call options “call -m -f gq”. SNPs and INDELs were annotated with SnpEff v5.0^104^. Several filters were applied to remove low-quality SNPs: average read depth across all samples < 10, maximum read depth > 128000, minimum mapping quality phred score < 40, minimum SNP quality phred score < 20, and multiallelic variants. DMS variants from our study were then matched with the identified genomic variants to determine the presence/absence and frequency of the mutations.

The visualization of protein structures was done with ChimeraX^105^. Figures and analysis were done using the following Python packages: Matplotlib^106^, NumPy^107^, UpSet^108^, SciPy^109^, pandas^110^ and seaborn^111^.

## Extended data

**Extended Data Fig. 1.**
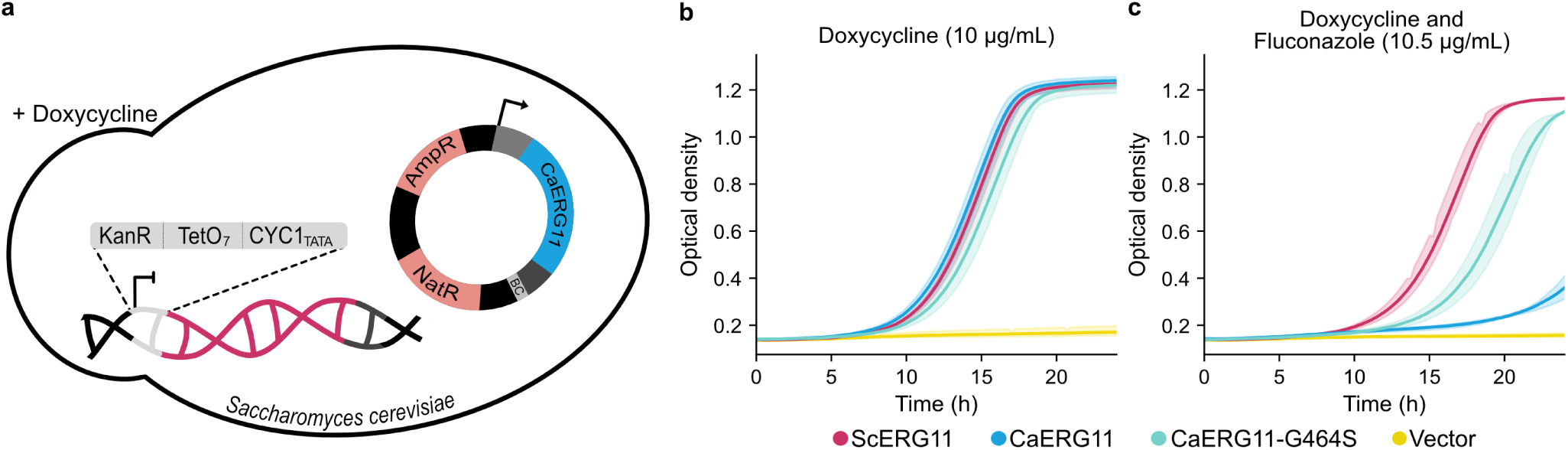
*ScERG11*-DOX strain repression and *CaERG11* complementation and resistance. **a,** The endogenous promoter of *ScERG11* was replaced with a doxycycline (DOX) repressible promoter that is fully activated in the absence of DOX, but repressed in the presence of DOX (strain *ScERG11*-DOX)^39^. *CaERG11* is expressed from a plasmid containing the native promoter and terminator of *ScERG11*, *CaERG11*, a 20N barcode (BC) and AmpR and NatR resistance markers. **b,c** OD_600_ was measured in YPD+NAT medium with DOX (10 μg/mL) to validate *ScERG11* repression in the *ScERG11*-DOX strain (**b**) and with DOX (10 μg/mL) and fluconazole (10.5 μg/mL) to validate *CaERG11* complementation and resistance (**c**). When *ScERG11*-DOX is transformed with an empty plasmid, there is no growth, as expected since *ERG11* is essential for *S. cerevisiae.* Growth is restored when *ScERG11* is expressed from the plasmid. Transformation with the *CaERG11* plasmid rescues growth, showing that *CaERG11* complements the function of *ScERG11*. *ScERG11*-DOX transformed with the *CaERG11* plasmid is susceptible to fluconazole. When *ScERG11*-DOX is transformed with the plasmid expressing *CaERG11*-G464S, a substitution known to confer fluconazole resistance^41^, growth has recovered. The colored bands represent standard deviations of the measurements, n=3.

**Extended Data Fig. 2.**
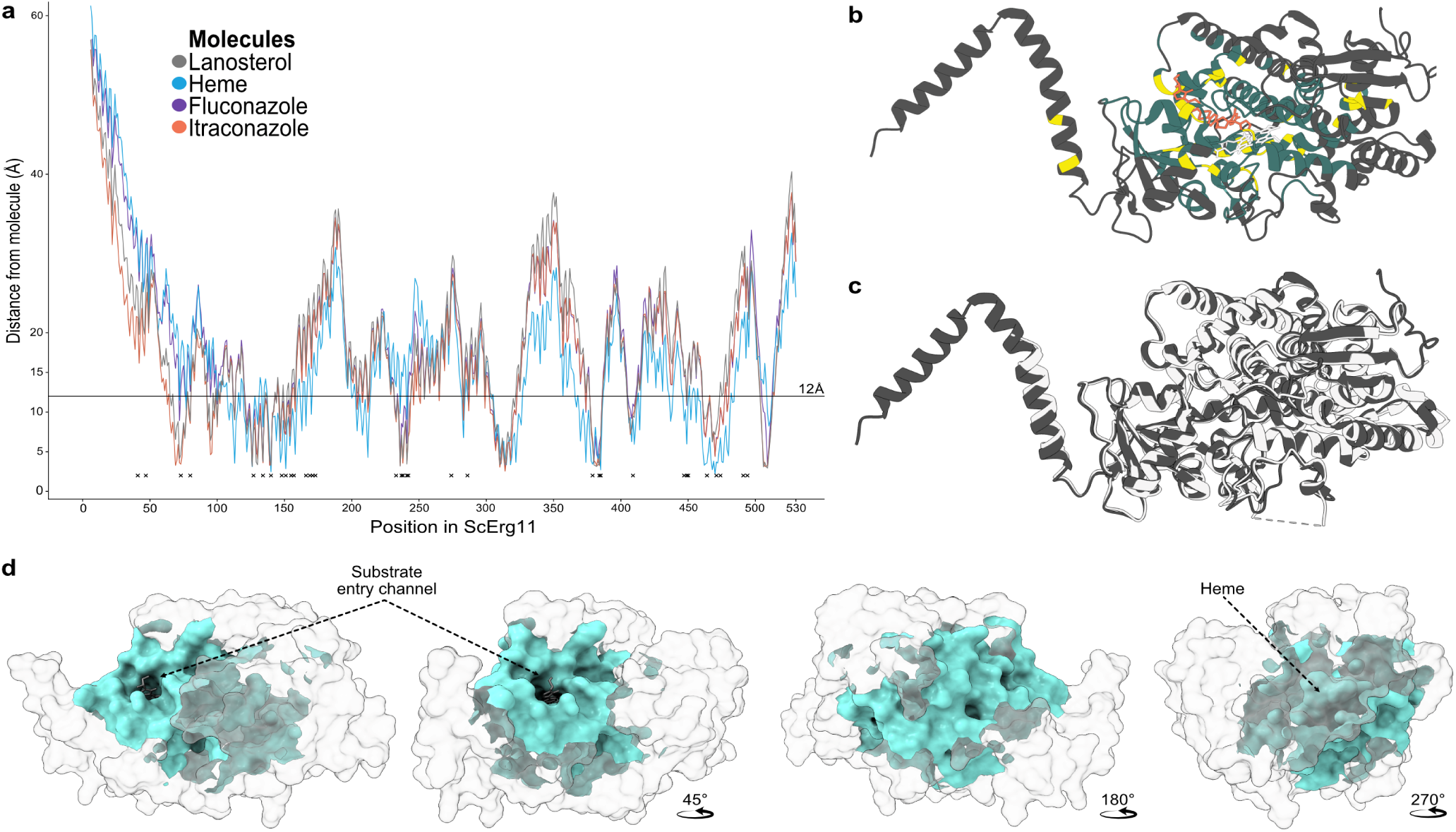
Selection of residues to be mutated in CaErg11 using the ScErg11 structures. **a,** Structures of ScErg11 bound to lanosterol (PDB ID: 4LXJ), fluconazole (PDB ID: 4WMZ) and itraconazole (PDB ID: 5EQB) were used to calculate the distances in ångström (Å) between the molecules and amino acids.The Xs represent positions with known resistance mutations listed in the MARDy database^84^. All residues at less than 12 Å from the lanosterol, the fluconazole, the itraconazole and the heme of ScErg11 were selected. **b,** We selected a radius covering most residues (206, dark green) with known mutations in the MARDy database (yellow^84^), even though most of them had not been validated. On the structure, ScErg11 is bound to itraconazole (orange) and an heme (white) (PDB ID: 5EQB). **c,** ScErg11 (dark grey, PDB ID: 5EQB) superposed on CaErg11 (light grey, PDB ID: 5V5Z). **d,** CaErg11 bound to itraconazole (PDB ID: 5V5Z). The 206 residues mutated in the variant library of CaErg11 surround the ligand binding site (cyan).

**Extended Data Fig. 3.**
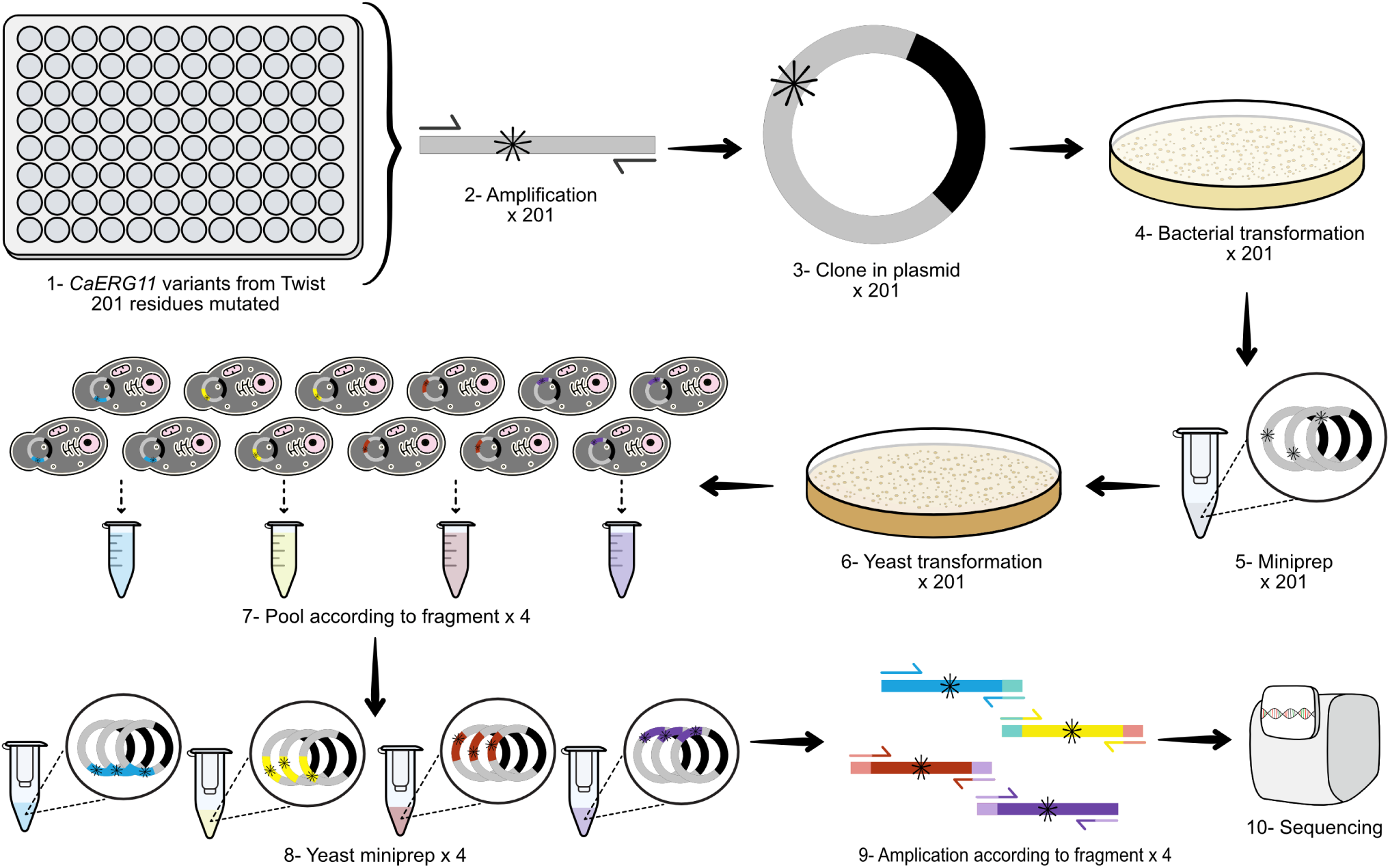
Construction and validation of the *CaERG11* variant library. 1- *CaERG11* DNA variants were synthesized (Twist Bioscience). Each well of 96-well plates contained DNA fragments corresponding to one mutated position. 2- *CaERG11* DNA variants (*) were amplified and 3- cloned in a plasmid containing the native promoter and terminator of *S. cerevisiae ERG11* to ensure that expression regulation is representative of the native gene. 4- Plasmids were transformed into bacteria, all colonies on the plates were scraped and 5- minipreps were done to retrieve the plasmids. 6- All the plasmids were transformed into the yeast strain *ScERG11*-DOX and colonies were collected from plates. 7- The *CaERG11* library was separated into pools corresponding to four overlapping fragments for screening. 8- Plasmids were extracted using yeast minipreps. 9- *ERG11* variants were amplified from plasmid DNA according to their corresponding fragments. 10- Variant diversity was assessed by high-throughput sequencing.

**Extended Data Fig. 4.**
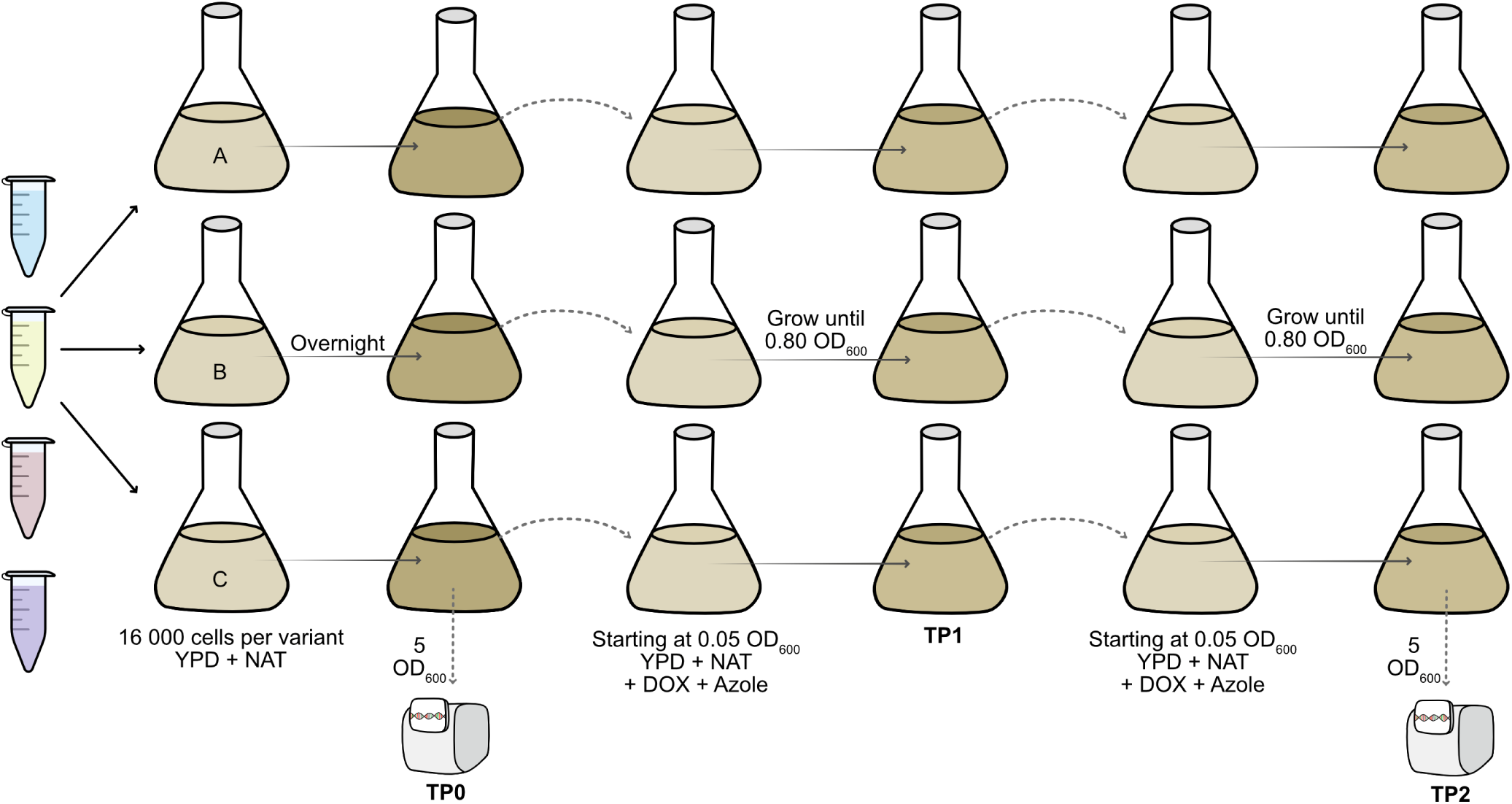
Pooled competition assay of CaErg11 variants. All fragment pools were grown overnight in three replicates (A, B and C) in 25 mL of YPD+NAT medium starting with around 16,000 cells per variant (Timepoint 0, TP0). The saturated cultures were transferred to fresh YPD+NAT with doxycycline and azole at a starting OD_600_ of 0.05 and grown until an OD_600_ of 0.80 was reached (Timepoint 1, TP1). The cultures were transferred again into fresh YPD+NAT with doxycycline and azole at a starting OD_600_ of 0.05, and grown until they reached an OD_600_ 0.80 (Timepoint 2, TP2). The average number of generations per transfer was four, for a total of eight generations. At the end of TP0 and TP2, 5 OD_600_ units of cells were used for plasmid extraction and sequencing.

**Extended Data Fig. 5.**
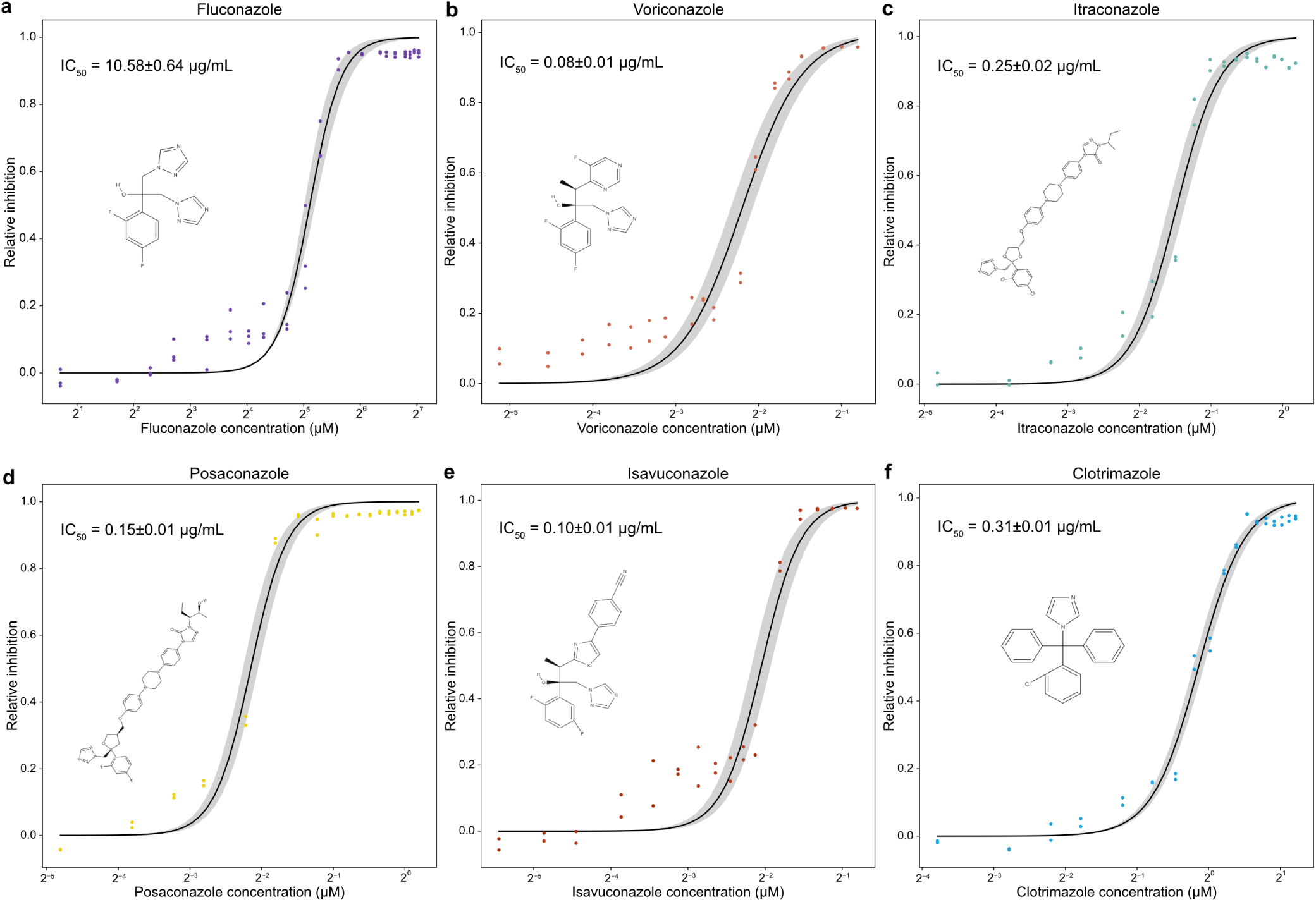
Dose-response curves for the engineered strain *ScERG11*-DOX in the presence of six medical azoles. **a,b,c,d,e,f,** Relative growth inhibition as a function of drug concentration of fluconazole (**a**), voriconazole (**b**), itraconazole (**c**), posaconazole (**d**), isavuconazole (**e**) and clotrimazole (**f**). The growth of the *ScERG11*-DOX strain expressing *CaERG11* from a plasmid was measured in rich YPD+NAT medium containing doxycycline (10 μg/mL). In each case, OD_600_ was measured every 15 minutes and the maximum growth rates were estimated from the growth curves. The maximum growth rates were transformed into relative inhibition. Each point here represents the relative inhibition at a given drug concentration. The concentration corresponding to 50% inhibition (IC_50_) is shown on the plots. Standard deviation is shown in light gray. Molecular structures were retrieved from PubChem^46^. n = 3 replicates for fluconazole and n = 2 for other drugs.

**Extended Data Fig. 6.**
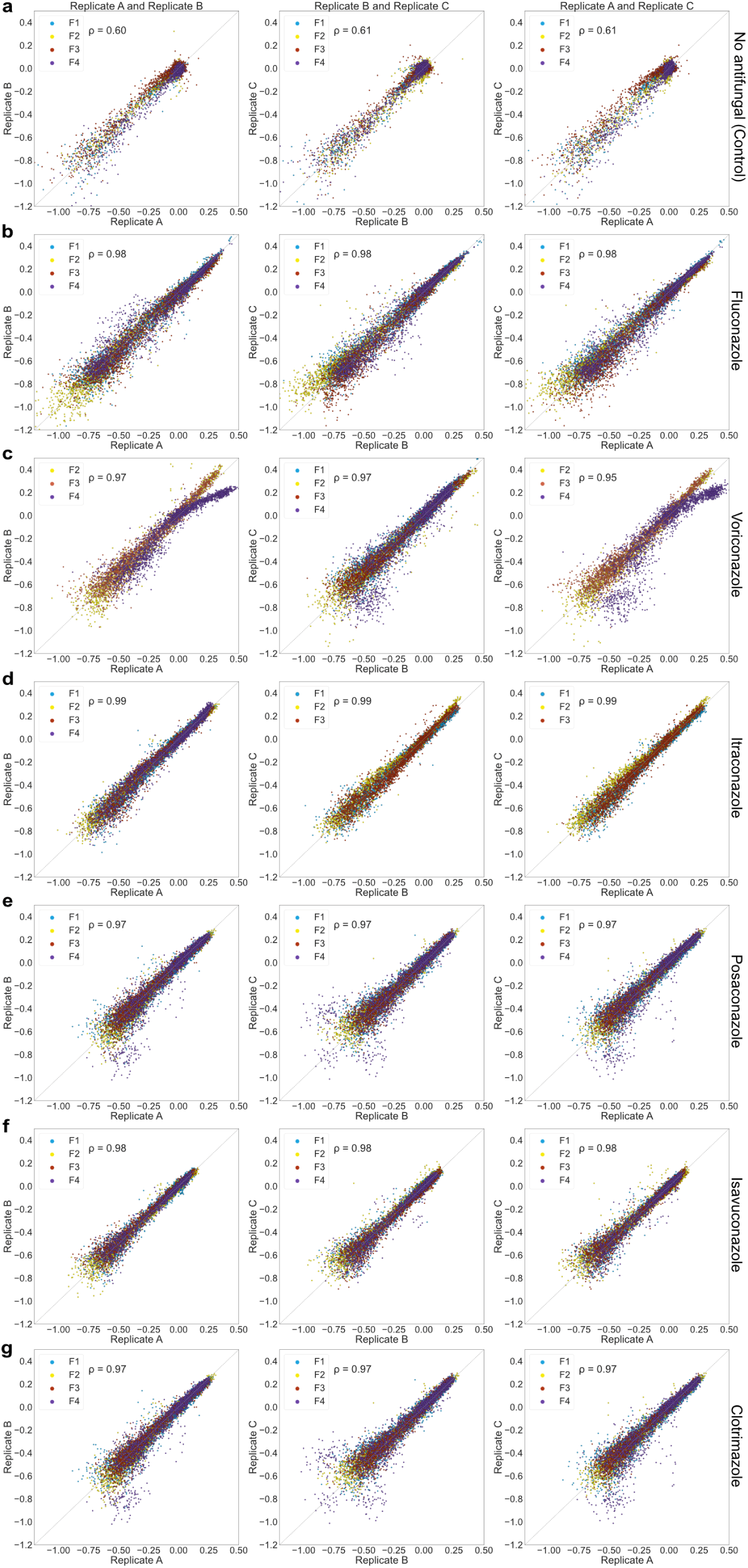
Correlation between estimates of selection coefficients for individual codon variants across replicated pooled competition assays. **a,b,c,d,e,f,g,**. In control (**a**) and in fluconazole (**b**), voriconazole (**c**), itraconazole (**d**), posaconazole (**e**), isavuconazole (**f**) and clotrimazole (**g**). Colors represent the four different fragments. Replicate A of F1 for voriconazole and Replicate C of F4 for itraconazole had to be removed since sequencing failed for these samples. Spearman’s rank correlation is shown, *P <* 0.0001 in all cases. n = 7,649 DNA variants.

**Extended Data Fig. 7.**
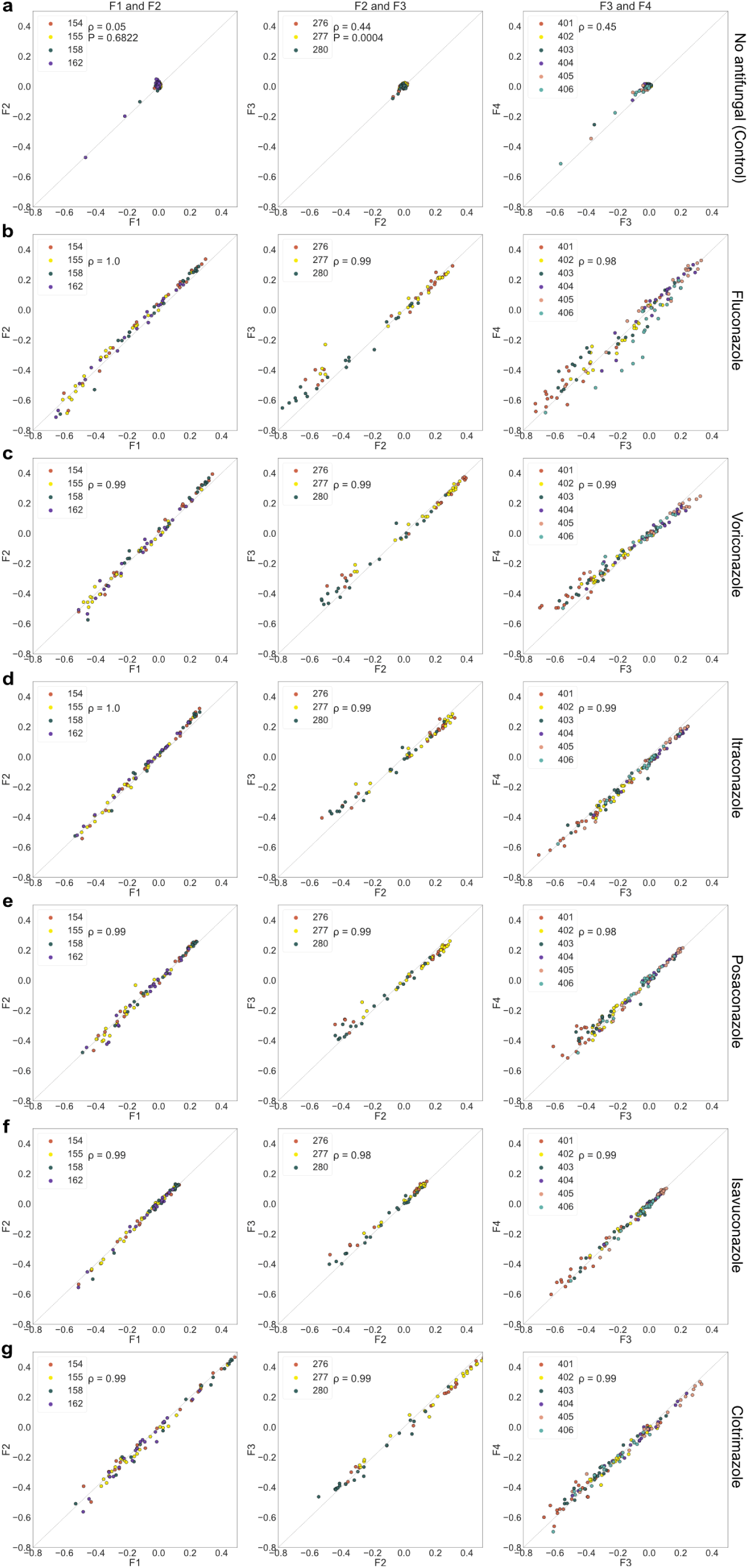
Correlation between overlapping fragment variants. **a,b,c,d,e,f,g,** The fragments of the library had overlapping sequences, which allowed us to assess the reproducibility of the assays by comparing the same variants in different pools of fragments. The selection coefficients are compared among fragments in control (**a**) and in fluconazole (**b**), voriconazole (**c**), itraconazole (**d**), posaconazole (**e**), isavuconazole (**f**) and clotrimazole (**g**). The values are the median selection coefficient of two synonymous codons per amino acid, and in most cases for the median of three replicates. Batch effect for the fragment 4 in the fluconazole condition was corrected with an interpolation model to get new F4 values following a linear regression with F3. Colors represent the different CaErg11 positions in the overlapping regions. Spearman’s rank correlation is shown, *P <* 0.0001 except when stated otherwise. F1-F2; n = 80 amino acid variants, F2-F3; n = 60 and F3-F4; n = 120.

**Extended Data Fig. 8.**
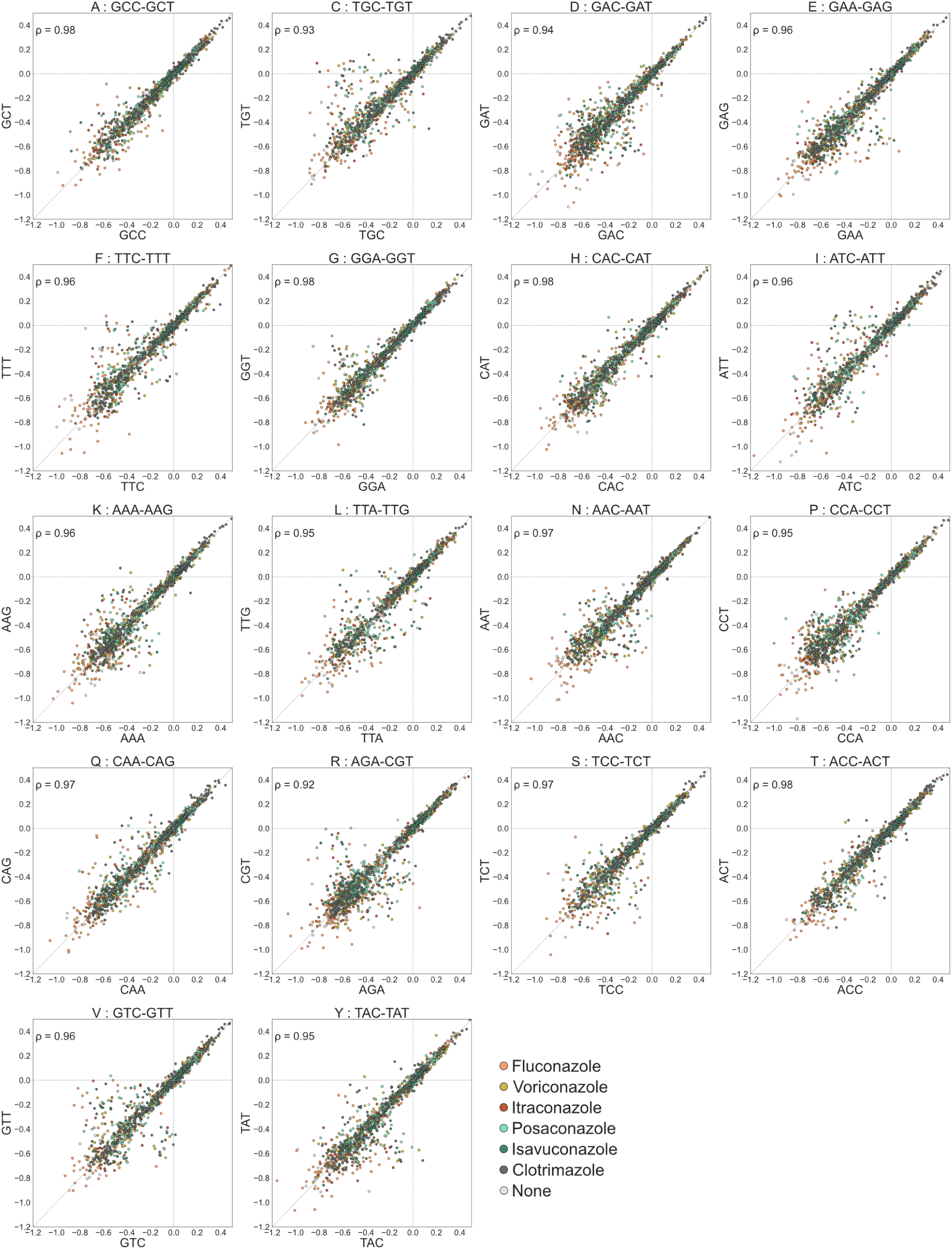
Correlation between synonymous codons. In the CaErg11 variant library, each of the 206 wild-type residues was replaced with the 19 non-wild type amino acids and this, using two codons when possible. To measure the reproducibility of the results, we looked at the correlation between the selection coefficients of synonymous codons for every amino acid (one letter abbreviation) for the median of three replicates. Colors represent the seven different pooled competition assays. Spearman’s rank correlation is shown, *P <* 0.0001. n = 7,448 DNA variants.

**Extended Data Fig. 9.**
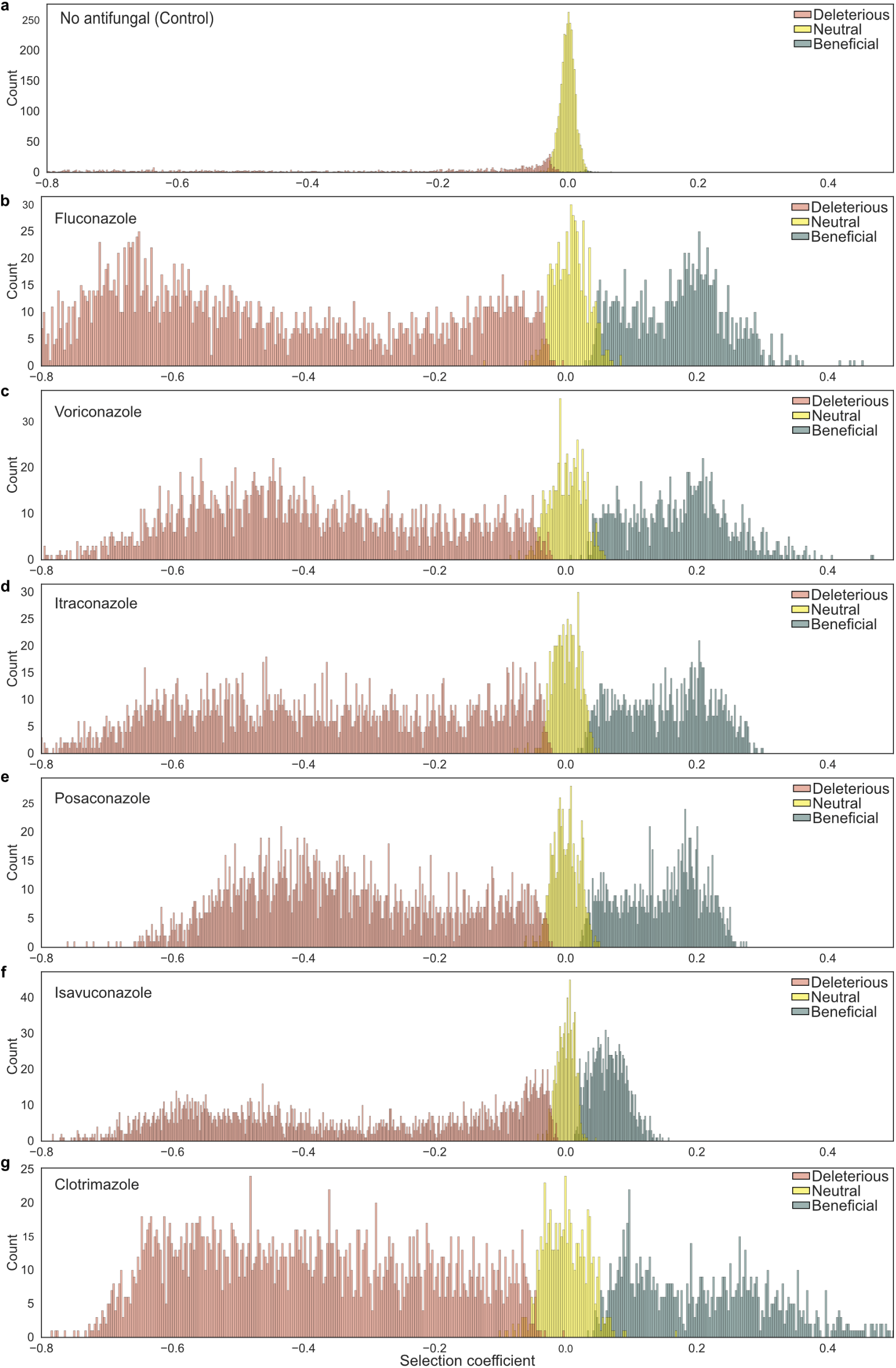
Distribution of Erg11 variant selection coefficients and their classification. **a,b,c,d,e,f,g,** Distribution of the selection coefficients in the control condition (**a**) and in fluconazole (**b**), voriconazole (**c**), itraconazole (**d**), posaconazole (**e**), isavuconazole (**f**) and clotrimazole (**g**). Colors represent the different categories. Categories were obtained by comparing the variants with wild-type synonymous codons to missense variants with two-sided independent *t-test* and by correcting *p-values* for multiple comparisons with the Benjamini-Hochberg method (deleterious: *s* < 0 and *P <* 0.01, neutral: *P >* 0.01, beneficial: *s* > 0 and *P <* 0.01). n = 3,830 amino acid variants.

**Extended Data Fig. 10.**
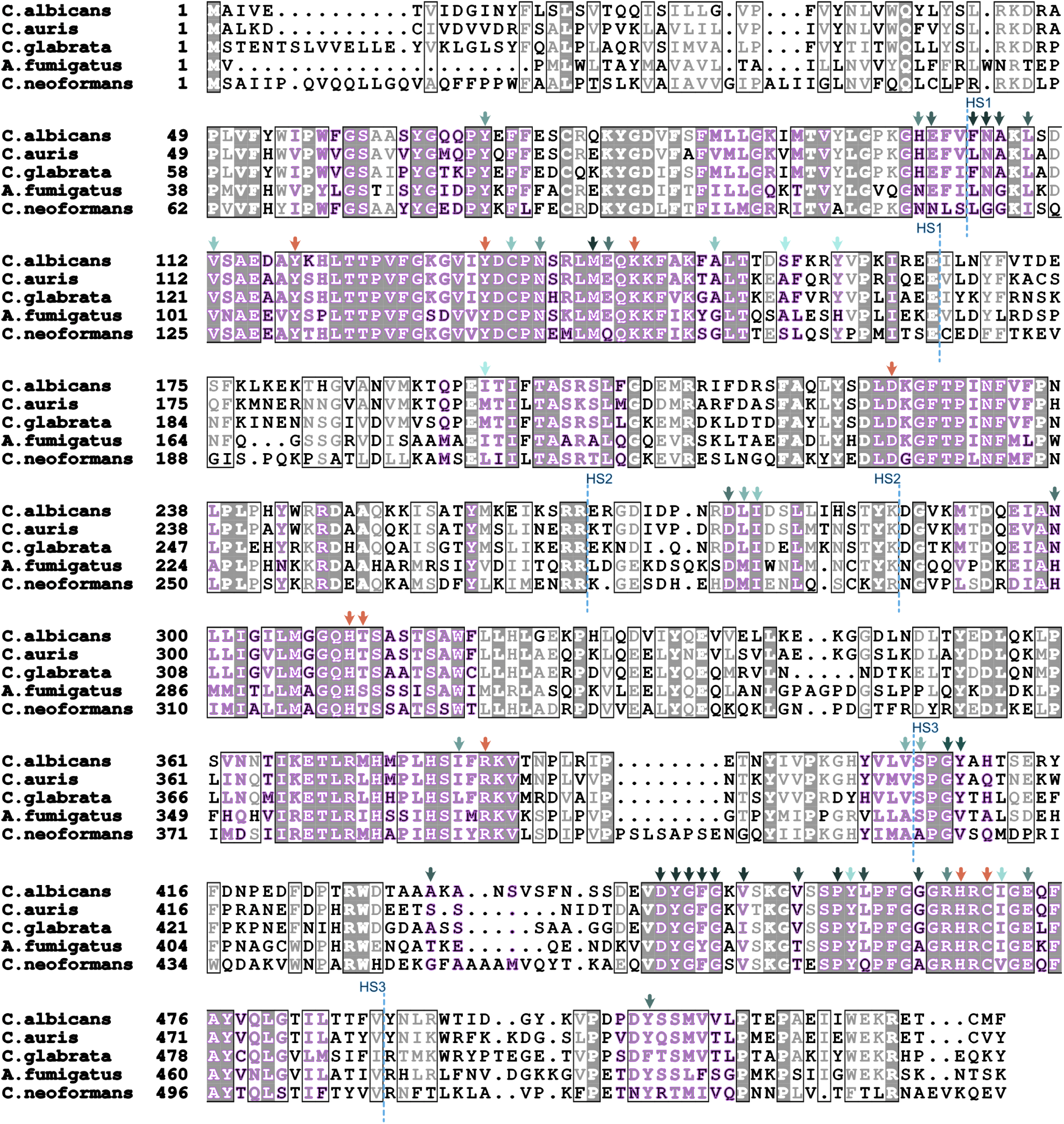
Multiple sequence alignment of Erg11 from some fungal pathogens. Amino acid positions outlined in purple were included in our CaErg11 library. The orange arrows point to amino acids interacting with the heme (Y118, Y132, K143, R381, H468 and C470) or participating in proton transport to the heme (D225, H310 and T311). Residues with more than 10 variants conferring fluconazole resistance are identified with green arrows (light green: 10/19 aa to dark green: 19/19 aa). They mostly fall into the three resistance clusters (HS) proposed by Marichal *et al*.^55^ (in blue, HS1 : 105 to 165, HS2 : 266 to 287 and HS3 : 405 to 488). The alignment was produced using ENDscript 2^54^.

**Extended Data Fig. 11.**
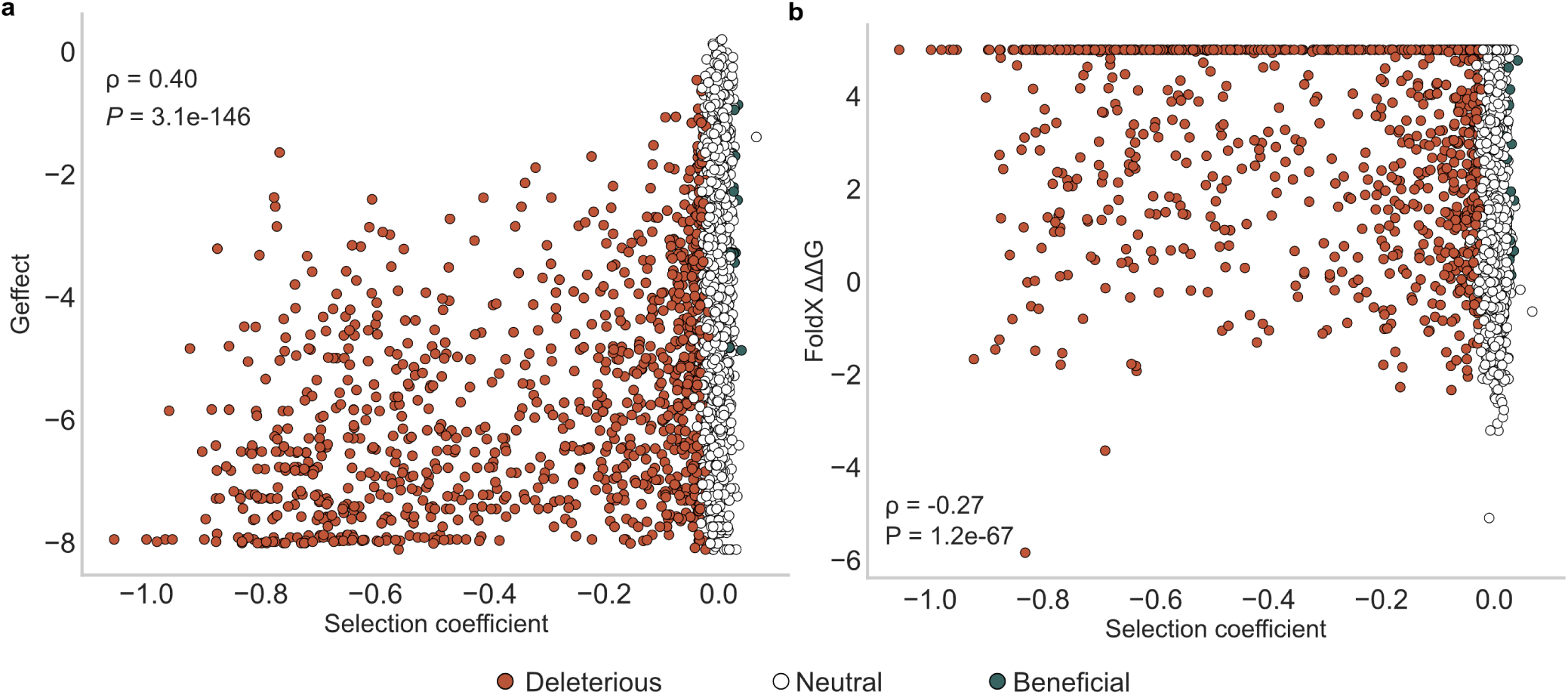
Estimation of the effect of mutations on the function of Erg11 using an evolutionary model (GEMME) and energy of protein folding modeling (FoldX). **a,** Relationship between the predicted impact of mutations inferred using long-term evolutionary constraints (Geffect) on Erg11 and selection coefficients in the control condition^48^. GEMME is a computational method that allows prediction of mutational outcomes by modeling the evolutionary history of sequences. Low Geffect values are predicted to be deleterious. Spearman’s rank correlation is shown. n = 3,830 amino acid variants. **b,** Examination of the relationship between the prediction of the effects of amino acid substitutions on CaErg11 (PDB ID : 5V5Z) stability from FoldX^49^ and the selection coefficient in the control condition. Positive ΔΔG values indicated destabilizing mutations. Spearman’s rank correlation is shown. n = 3,830 amino acid variants. The values above ΔΔG of 5 are rounded to 5.

**Extended data Fig. 12.**
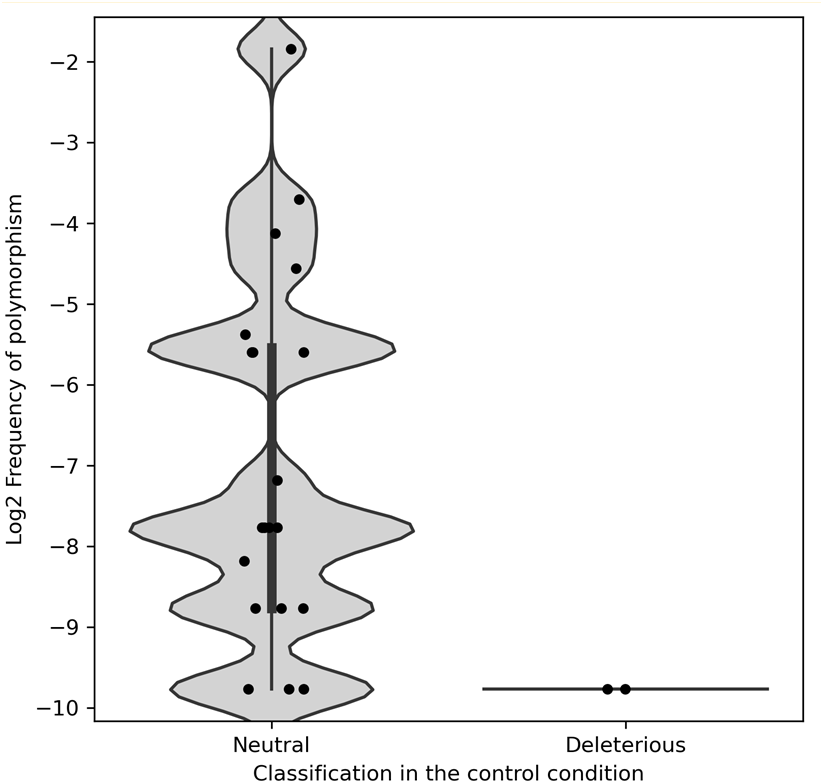
Frequency of polymorphisms in Erg11 of 435 *Candida albicans* genomes. Frequency of each polymorphism in Erg11 detected among the 435 genomes and present in our DMS library in Log2. The amino acid polymorphisms are separated based on their classification of fitness effect from our assays in the control condition. Mutations we classified as deleterious in the control condition are much less frequent than the ones we classified as neutral in *C. albicans* clinical isolates.

**Extended Data Fig. 13.**
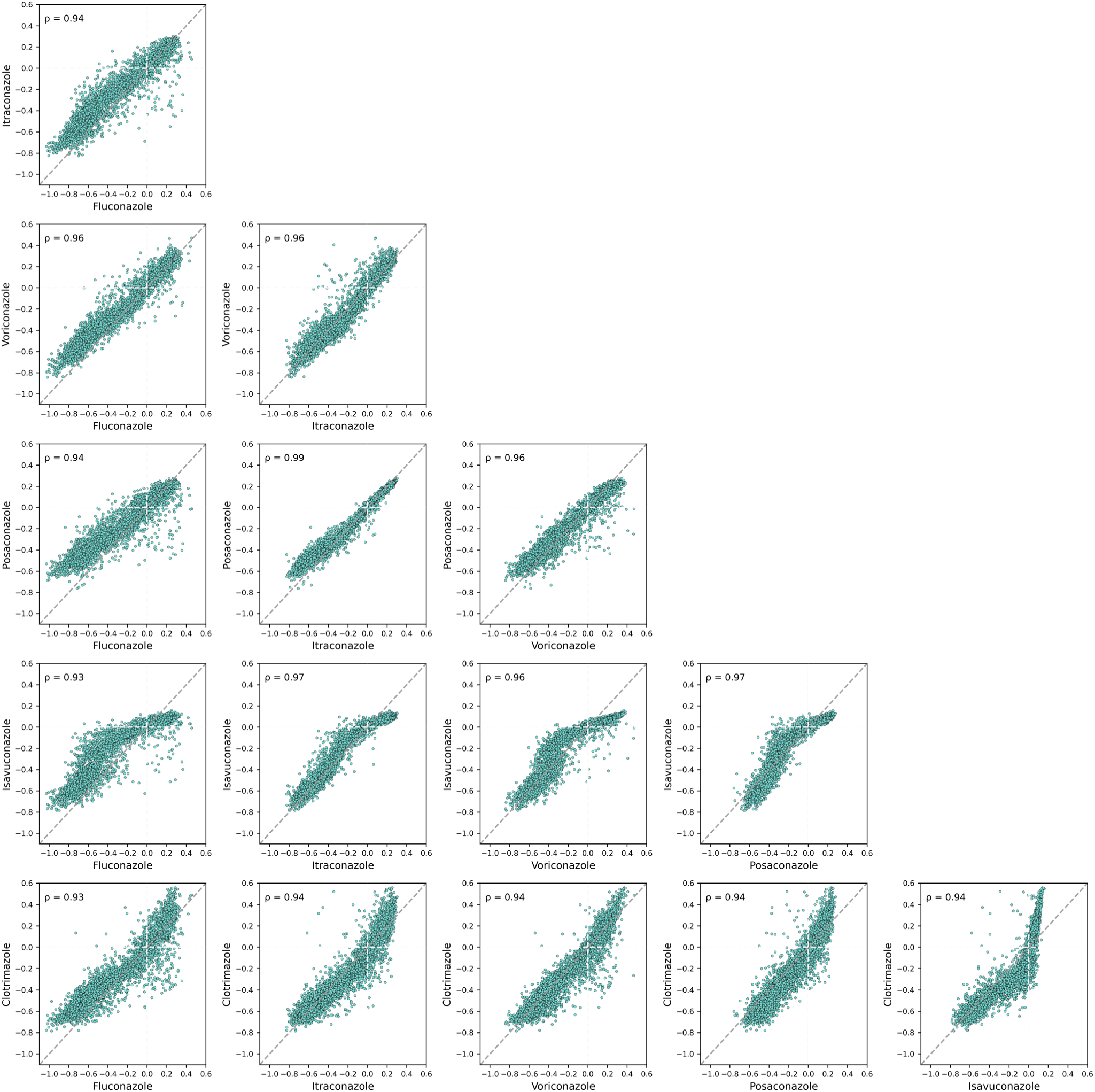
Correlation between Erg11 variant selection coefficients in the presence of different azoles. Each value is the median of two synonymous codons per amino acid in most cases, and the median of triplicate experimental measurements. Spearman’s rank correlation is shown, *P <* 0.0001. n = 3,830 amino acid variants.

**Extended Data Fig. 14.**
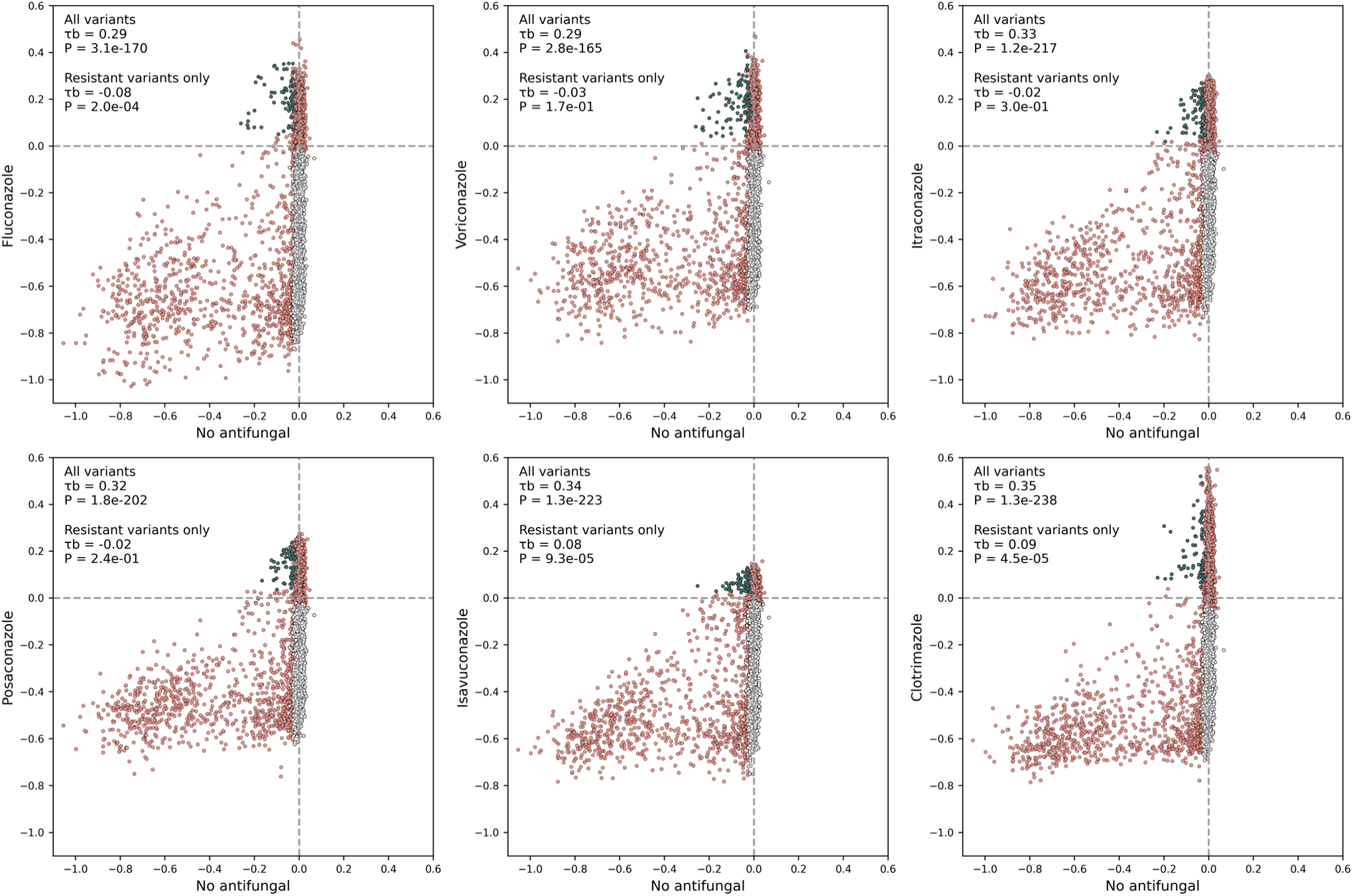
Correlation between Erg11 variant selection coefficients estimated in azole and in control conditions. Each value is the median of two synonymous codons per amino acid in most cases for the median of triplicate experimental measurements. The dark green dots represent variants with a significant tradeoff, meaning that the mutation is significantly advantageous in azole and deleterious in the control condition. The light grey dots represent deleterious variants in the presence of an azole but not in its absence. Kendall’s tau-b (τb) rank correlation was calculated for all variants and for the subset of resistant variants only. n = 3,830 amino acid variants.

**Extended data Fig. 15.**
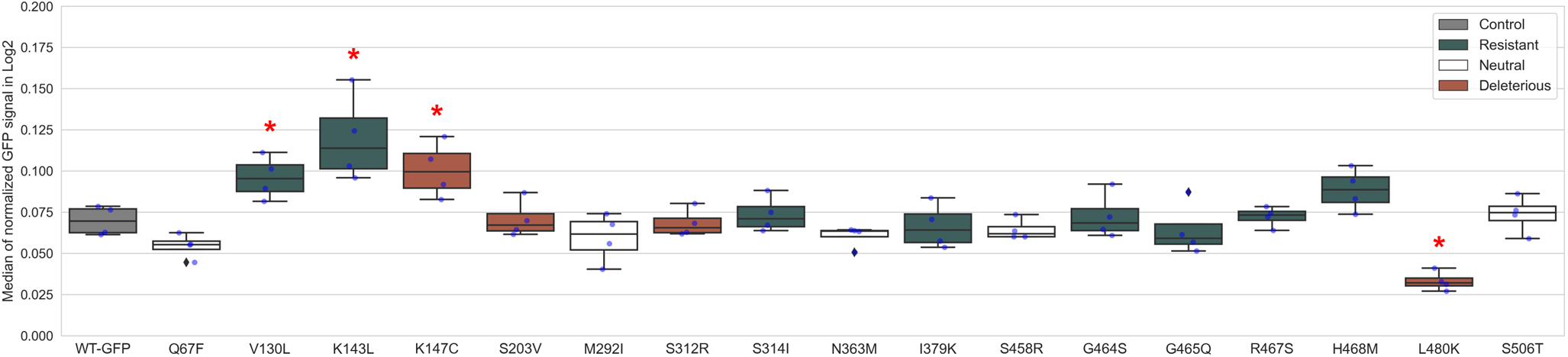
Quantification of CaErg11 abundance by flow cytometry. Median of the normalized GFP fluorescence signal in Log2 of *ScERG11*-DOX cells expressing CaErg11 tagged with the GFP (n = 17 mutants and n=1 wild-type). Significant difference with the wild-type (WT-GFP) is indicated by a red * (Mann-Whitney U test, *P* < 0.05). Colors represent the categorization of each mutant in the presence of fluconazole. n = 4 biological replicates.

## Notes

https://github.com/Landrylab/Bedard_et_al_2023

### Competing Interest Statement

The authors have declared no competing interest.

### Summary of Updates

One figure has been split, some more experiments were added.

